# PLSKO: a robust knockoff generator to control false discovery rate in omics variable selection

**DOI:** 10.1101/2024.08.06.606935

**Authors:** Guannan Yang, Ellen Menkhorst, Evdokia Dimitriadis, Kim-Anh Lê Cao

**Affiliations:** Melbourne Integrative Genomics, School of Mathematics and Statistics, The University of Melbourne, Parkville, Victoria, Australia; Department of Obstetrics and Gynaecology, School of Mathematics and Statistics, The University of Melbourne, Parkville, Victoria, Australia; Gynaecology Research Centre, School of Mathematics and Statistics, The University of Melbourne, Parkville, Victoria, Australia

## Abstract

The knockoff framework, combined with variable selection procedure, controls false discovery rate (FDR) without the need for calculating *p*−values. Hence, it presents an attractive alternative to differential expression analysis of high-throughput biological data. However, current knockoff variable generators make strong assumptions or insufficient approximations that lead to FDR inflation when applied to biological data.

We propose Partial Least Squares Knockoff (PLSKO), an efficient and assumption-free knockoff generator that is robust to varying types of biological omics data. We compare PLSKO with a wide range of existing methods. In simulation studies, we show that PLSKO is the only method that controls FDR with sufficient statistical power in complex non-linear cases. In semi-simulation studies based on real data, we show that PLSKO generates valid knockoff variables for different types of biological data, including RNA-seq, proteomics, metabolomics and microbiome. In preeclampsia multi-omics case studies, we combined PLSKO with Aggregation Knockoff to address the randomness of knockoffs and improve power, and show that our method is able to select variables that are biologically relevant.

## 1 Introduction

High-throughput technologies are widely used to measure system-wide biological variables, such as genes, proteins, metabolites, bacteria. Analytical methods have been developed to extract useful information from these datasets. For example, differential expression (DE) analysis aims to identify biological variables whose expression or abundance levels vary between experimental groups; penalised regression aims to identify variables that are predictive of an outcome variable. These variables are called ‘discoveries’, and can then be used to generate hypotheses for further validation and investigation. It is therefore important to generate reliable discoveries that are not false positives.

False discovery rate (FDR) is a statistical criterion to control false positives in discoveries. Controlling FDR under a certain cutoff (e.g., 0.05) ensures the reliability of multiple testing in high-throughput data analysis. Most FDR control approaches, such as the Benjamini-Hochberg (BH) procedure rely on a set of valid *p*-values (Benjamini and Hochberg, 1995). However, the calculation of *p*-values requires strong distribution assumptions and is only achievable under a simple model or a simple estimation. For more complex data such as high-throughput data, valid *p*-values are difficult to calculate.

Barber and Candès (2015) introduced ‘knockoffs’, a novel framework for obtaining FDR control in feature selection while bypassing the standard calculation of *p*-values. The basic principle of the knockoff procedure is to create artificial ‘knockoff’ variables that resemble the true correlation structure of the original data, but without using any information from the response variable. The original variables and their knockoff copy are run together into a model to measure the importance of each variable, so that the knockoff variables act as negative controls. A threshold is then derived to select the important variable set with controlled FDR.

The knockoff filtering procedure can be used with any feature selection method that generates importance measures under certain conditions. The aim is to identify the true important variables that are directly related to response of interest, while excluding variables that are indirectly related to the response, but are correlated with the true important ones. However, the application of knockoff to high-throughput biological data remains limited. The procedure heavily depends on the construction of knockoff variables, which can be challenging in practice when the distributions of the explanatory variables are unknown. Biological data exhibits remarkable diversity, encompassing a wide range of data types derived from measuring various biological molecules or entities, and from various technological platforms. Even when considering the marginal distribution of a single biological variable, assumptions on the data distribution can be questionable (Li et al., 2022). A second-order knockoff procedure was introduced assuming variables follow a multivariate Gaussian distribution (Candès et al., 2018), but this can over-simplify the distribution of biological data. Furthermore, the joint distribution of biological variables is complex given their dependency structure (e.g RNA sequencing, microbiome), thus invalidating the assumptions of second-order knockoffs.

Besides the distribution assumption of the knockoff procedures, another challenge is the high-dimensionality of the data, where the number of variables far exceeds the number of samples. The ‘curse of dimensionality’ leads to either underfitting or overfitting modelling. For example, KnockoffScreen (He et al., 2021) can underfit on the conditional distribution when the number of samples is small, leading to inflated FDR as it fails to control on other variables. Another example is second-order-approximation knockoffs based on sample covariance, which is prone to overfitting and generates knockoff variables identical to the original ones, resulting in a lack of power (Candès et al., 2018). Therefore, knockoff variable generators need to be robust to different types of distributions, and be able to fit high-dimensional biological data efficiently.

We developed a new knockoff variable generator, Partial Least Squares Knockoff (PLSKO) as a first step of knockoff filtering. PLSKO is efficient, assumption-free and applicable to various types of data distribution, such as omics data. We demonstrate the robustness and improved performances in FDR control and statistical power compared to a wide range of existing knockoff generators in both simulation and semi-simulation studies. We apply PLSKO to multi-omics studies in preeclampsia, and compare our results to classical differential analysis. The selected features can be used to identify biomarkers and for downstream analyses and modelling.

## 2 Methods

### 2.1 Knockoff filtering: FDR control in variable selection

In the following, we denote *y* a clinical outcome or phenotype variable of interest with *n* observations, and **X** an (*n* × *p*) expression matrix where *p* is the number of omics variables (e.g., genes, proteins). *X*_*j*_ represents a column from **X** for gene *j*. We first introduce knockoff filtering then describe our proposed approach PLSKO.

#### Controlled variable selection

Knockoff filtering is a procedure for controlled variable selection. Knockoff aims to select ‘relevant’ variables that are conditionally dependent on the response variable given all the other covariates, that is, a subset of **X** that directly affects *y. X*_*j*_ is referred to as ‘null’ if it is conditionally independent of *y* given the other *p* − 1 variables. As we consider the correlation among the variables into account, controlled variable selection ensures the hypotheses are mutually independent in multiple testing, thus avoiding ill-posed type-I error control and improving power. In practice, controlled variable selection improves interpretability of the results and makes further investigation of causal relationships possible.

#### False Discovery Rate (FDR)

Given any variable selection procedure, a selected variable set is an estimate of the true important variable with random error. FDR is the expected false discovery proportion (FDP), that is, the proportion of null variables among all selected variables. FDP fluctuates around the FDR in any FDR-controlling procedure due to the randomness in both data and selection procedure. A knockoff filter selects variables controlling the FDR with finite sample guarantees.

### Main steps of the knockoff filtering

Knockoff filtering consists of three main steps (summarised in Supplemental Figure S1 and described in details in Section S3.1):

1. **Knockoff variable construction**. Knockoff variables, 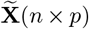, can be generated satisfying two properties of model-X knockoff for high-dimensional data where *p* > *n* (Candès et al., 2018): (1) the joint distribution of the original variables and the knockoff variables remain the same when swapping any subset of variables with their knockoff; and (2) the knockoff variables are constructed regardless of the information from *y*. As a ‘negative control’, a knockoff variable contains all the information from its original variable, and this information can be reconstructed by the other variables. We describe existing approaches to construct model-X knockoff in Supplemental Method S3.2.
2. **Important statistics calculation**. A variable selection model (e.g. lasso penalised regression) is then run on *both* original and generated knockoff variables as covariates, with the response variable *y*. Importance scores *W*_*j*_, *j* = 1, …, *p*, are obtained for *X*_*j*_ as ‘true’ importance score that remove the ‘negative control’ part under the ‘flip-sign’ property. A large positive value of *W*_*j*_ provides some evidence that the distribution of *y* depends upon *X*_*j*_, whereas small positive or negative values provide evidence of null variables. For example, an option to calculate *W*_*j*_ is the lasso coefficient difference (LCD) between *X*_*j*_ and its knockoff 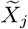.
3. **Threshold identification and variable selection**. To control FDR at a pre-specified level *q* (e.g. 0.05), we determine the threshold *T* as the minimum value that allows the proportion of variables with negative *W* ≤ −*T* compared to the variables with positive *W* ≥ *T* lower (or equal) than *q*. With a well-defined *W*, the number of null variables in the selected variable set can be estimated by the number of variables with large negative *W* which is expected to be equal to the number of null variables (i.e., false discoveries with large positive *W*). We select variables with *W*_*j*_ ≥ *T* with modified FDR controlled. Alternatively, we can increment the number of negatives in discoveries to identify the threshold *T*_+_ to control FDR in a slightly conservative manner (both options are detailed in Supplemental Method S3.1).

#### Sequential Conditional Independent Pairs (SCIP)

SCIP has been proposed as a general algorithm for exact knockoff variable construction. It does not require distribution assumptions on **X** (Candès et al., 2018). SCIP forms the basis of our proposed approach. The algorithm (described in Supplemental Algorithm S1) is based on the pairwise exchangeability property of model-X knockoffs, that is, a pair of a variable and its knockoff is distributionally non-distinguishable conditioning on all other variables and their knockoffs. To reduce computational costs in this sequential algorithm, we can generate 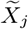 by sampling from the conditional distribution of *X*_*j*_ given the neighbours of *X*_*j*_ and 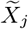 with the assumption of conditional independence to non-neighbours (He et al., 2021).

### 2.2 Knockoff variable construction for biological data

Our new method, ‘PLSKO’ incorporates partial least squares (PLS, Wold 1966) regression into the model-X knockoff framework. PLSKO overcomes the limitations of existing knockoff generators of biological data, which tend to either under- or over-fit due to high-dimensionality and unspecific distribution.

#### Partial Least Squares regression (PLS)

PLS encompasses a wide class of techniques for modelling relations between sets of variables. The assumption of PLS is that predictors and the response are driven by a small number of latent variables. PLS has been widely applied to high-dimensional biological data due to its high computational and statistical efficiency when *p* > *n* (Boulesteix and Strimmer, 2007). In our proposed PLSKO, we use PLS regression to calculate the conditional distribution in a sequential manner, with the aim of effectively controlling the information explainable by other variables.

#### Main steps of PLSKO

Given a set of biological variables **X**, we first need define neighbour variables, such as variables highly correlated with with *X*_*j*_, or a pre-defined list (e.g. genes from a specific pathway). By assuming a variable is only dependent on its neighbours and is independent with its non-neighbours, PLSKO reduces the number of variables being controlled in the conditional distribution, thus boosting computational time.

PLSKO generates knockoff variables from the first variable in the dataset to the last, individually, as in the SCIP algorithm (see Figure 1). For each variable *X*_*j*_, *j* = 1… *p*, PLSKO calculates the conditional distribution of *X*_*j*_ by fitting a PLS regression where *X*_*j*_ is the response and the covariates are its neighbour variables and neighbours’ constructed knockoff variables. The sample residuals from the regression can be considered as observations from the conditional distribution. By sampling from the estimated conditional distribution, we then permute the sample residuals. Each knockoff variable 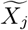 is then generated by adding the fitted value and permuted residual. This permutation-based is non-parametric and ensures PLSKO can be robustly applied to different types of data with unspecified distribution while controlling FDR. From PLSKO, we obtain the knockoff variable dataset 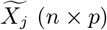 as the output of the first step of knockoff filtering. We can then proceed with the subsequent two steps of the knockoffs described above to select important variables. The full methodological details are available in Supplemental Methods S3.3 and Algorithm S2.

**Figure 1:**
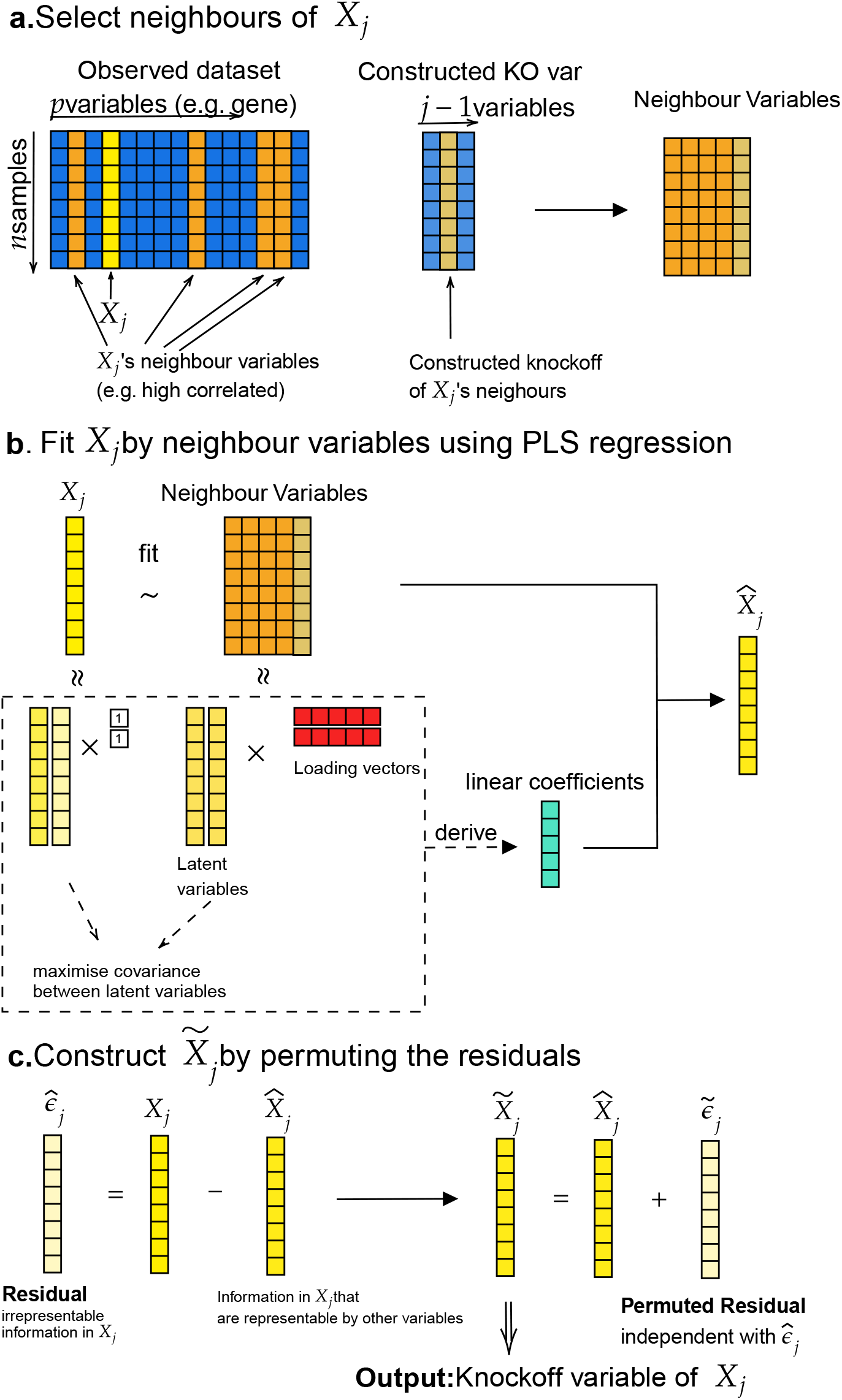
Steps of PLSKO to generate a knockoff variable 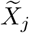 of *X*_*j*_. a. We define neighbour variables that are either highly correlated with *X*_*j*_, or a pre-defined list; b. We then fit a PLS regression for *X*_*j*_ using its neighbours and their knockoffs as predictors; c. Finally, we calculate then permute the residuals to create the knockoff 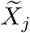. We iterate this process for each of the variables *X*_*j*_, *j* = 1,*…, p*.

#### Parameter specification of PLSKO

There are three types of parameters to consider in PLSKO:

- *Neighbour list*. In our default setting, we define neighbours based on sample correlations. The threshold value can be pre-specified, or based on a quantile of all correlations, depending on how sparse the co-expression network is assumed to be. However, the lower the threshold of the neighbourhood, the more similar the knockoff becomes to the original variables. This increased similarity can potentially lead to overfitting, reduced power, and a smaller Type I error. Therefore, a trade-off needs to be achieved.
- *Number of latent components in PLS*. The larger the number of latent components, the more similar the knockoff to the original variables, which can lead to a smaller type I error but also smaller power.
- *Variable kept in sparse PLS*. During the conditional distribution calculation (Step b. in Figure 1), we provide the option of sparse PLS (sPLS Lê Cao et al. 2008) to improve model fitting. The percentage of variables kept on each component is user-defined.

#### PLS-AKO

In our case studies, we used the Aggregation of Multiple Knockoffs (AKO) procedure to improve stability and reduce randomness in knockoff generation (Nguyen et al., 2020). We refer to this method as ‘PLS-AKO’. We generated multiple knockoff datasets using PLSKO (e.g. 50 in this study), then calculated LCD as the importance score, based on which we computed an empirical *p*-value for each *X*_*j*_ in each run. These empirical *p*-values were aggregated using a quantile-based method for each *X*_*j*_. Finally, we controlled the FDR by applying the BH procedure to the aggregated *p*-values, selecting variables accordingly. This approach generally resulted in a set of selected variables with lower FDR, higher power and stability compared to the average of single-run knockoffs. Details can be found in Supplemental Method S3.3.1.

### 2.3 Benchmark methods

We performed a series of simulation experiments with varying settings to compare the performance of knockoff generators, including second-order approximation methods, SCIP methods and PLSKO. We calculated the false discovery proportion (FDP) and true positive proportion (TPP) to evaluate their performance.

We considered second-order approximation (SOA) methods including semidefinite program (SDP) knockoff (Candès et al., 2018)) and minimised reconstructability (MRC) knockoff with both minimum variance-based reconstructability loss (MVR (Spector and Janson, 2022)) and maximised entropy loss (ME knockoff Gimenez and Zou 2019). Since this type of methods requires mean and covariance of **X** to be known or estimated, our benchmark considered ground truth covariance when available (namely ‘oracle’) or James-Stein (JS) style shrunk covariance (Ledoit and Wolf, 2003), the default method in the `knockoff` R package for high-dimensional data. SCIP-based knockoff generators include `SeqKnockoff` (seqko) with lasso regression for conditional distribution approximation (Kormaksson et al., 2021) and `KOBT` with PC regression (PCKO, Jiang et al. 2021). We also evaluated a knockoff generator for data generated from a factor model, namely ‘Intertwined probabilistic factors decoupling’ (`IPAD`) (Fan et al., 2020). We detail these methods in Supplemental Method S3.4 and S3.5.

In our benchmark, we examined the effect of various neighbourhood thresholds and the number of components in PLSKO, denoted as:

- PLSKO: Arbitrary and liberal setting: neighbour threshold of 80% quantiles (i.e. the top 20% of the most correlated pairs are used for PLS fitting) and 5 components.
- PLSKO-full: Most conservative setting. No neighbour filtering and all other variables are used for fitting. In simulations, the number of factors is set to be the known true rank of the whole data; for real data, the number of factors of PLSKO is chosen by the *PC*_*p*1_ criterion from Bai and Ng (2002).
- PLSKO-full-sparse: Parameters are the same as PLSKO-full but using sPLS regression with 20% of variables selected.

### 2.4 Preeclampsia studies

Preeclampsia is a life-threatening disease of pregnancy and a leading cause of maternal and neonatal morbidity and mortality, diagnosed by sudden-onset hypertension (> 20 weeks of gestation) and other maternal organ or placental dysfunction (Dimitriadis et al., 2023). We analysed two case-control studies of preeclampsia for both our semi-simulations and our case studies. The first study contains maternal circulating cell-free RNA-seq (cfRNA) data from 71 samples (16 preeclamptic, 55 normotensive) (Moufarrej et al., 2022). The second study is a multi-omics study that includes proteomics, metabolomics and microbiome data from 36 samples (cohort 1, 18 preeclampsia, 18 normotensive) or 49 samples (cohorts 1 and 2 combined for microbiome, 29 preeclamptic, 20 normotensive) (Marić et al., 2022). To avoid repeated measurements and maintain the assumption of independent samples in knockoff filtering, we considered only samples collected before 12 weeks of gestation (corresponding to the first sampling time) in our analysis.

## 3 Simulation Studies

Our simulations aimed to answer the following: (1) Does PLSKO control FDR with high power? and, (2) How robust are knockoff generators when their assumptions are violated?

### Simulation setting

Most knockoff generators have been shown to control FDR in simplistic simulations, where **X** follow a multivariate Gaussian distribution. We have confirmed these results, and also showed that the FDR was controlled when **X** is generated from a Gaussian factor model (Supplemental Figures S2, S3, S4 and S5). In these scenarios, the relationships between variables are linear, which may not capture the complexity of real biological data. To test whether PLSKO and other existing methods can robustly handle more challenging cases such as when the relationships between variables are non-linear, we simulated data from a quadratic factor model: half of the variables were generated from a block factor model and the other half as their square value. We randomly selected a subset of variables known as important variables to generate a response *y* as a linear combination of these variables. For each given parameter configuration, we simulated 100 high-dimensional datasets, with *p* = 500. We then generated knockoff variables with different knockoff generators, and used the lasso coefficient difference (LCD) as the importance statistics *W* in knockoff filtering with a target FDR threshold *q* = 0.05. The FDP and the true positive proportion (TPP) of the selected variable set were used to assess the validity and performance of the knockoff generators. The average FDP gives an estimate of FDR, whereas the average TPP gives an estimate of power. Our simulations are detailed in Supplemental Method S3.5 and Table S5.

### Simulation results

PLSKO with 12 components was the only method that empirically controlled the FDR for any sample size, while PLSKO with 9 components and sequential knockoff (seqko) had relatively low FDR *<* 0.10 (Figure 2). Other methods had a large FDR. Especially, the factor-model-based methods IPAD and PCKO had an FDR > 0.15, even for a number of components in the Principal Component regression similar to those in PLSKO (9 and 12). Due to its iterative nature, the PLS regression results in a better fit in controlling other variables with a non-linear relationship compared to Principal Component regression in PCKO. PLSKO was highly efficient as it utilises a relatively small number of components, and its power of PLSKO did not reduce when we increased the number of components.

**Figure 2:**
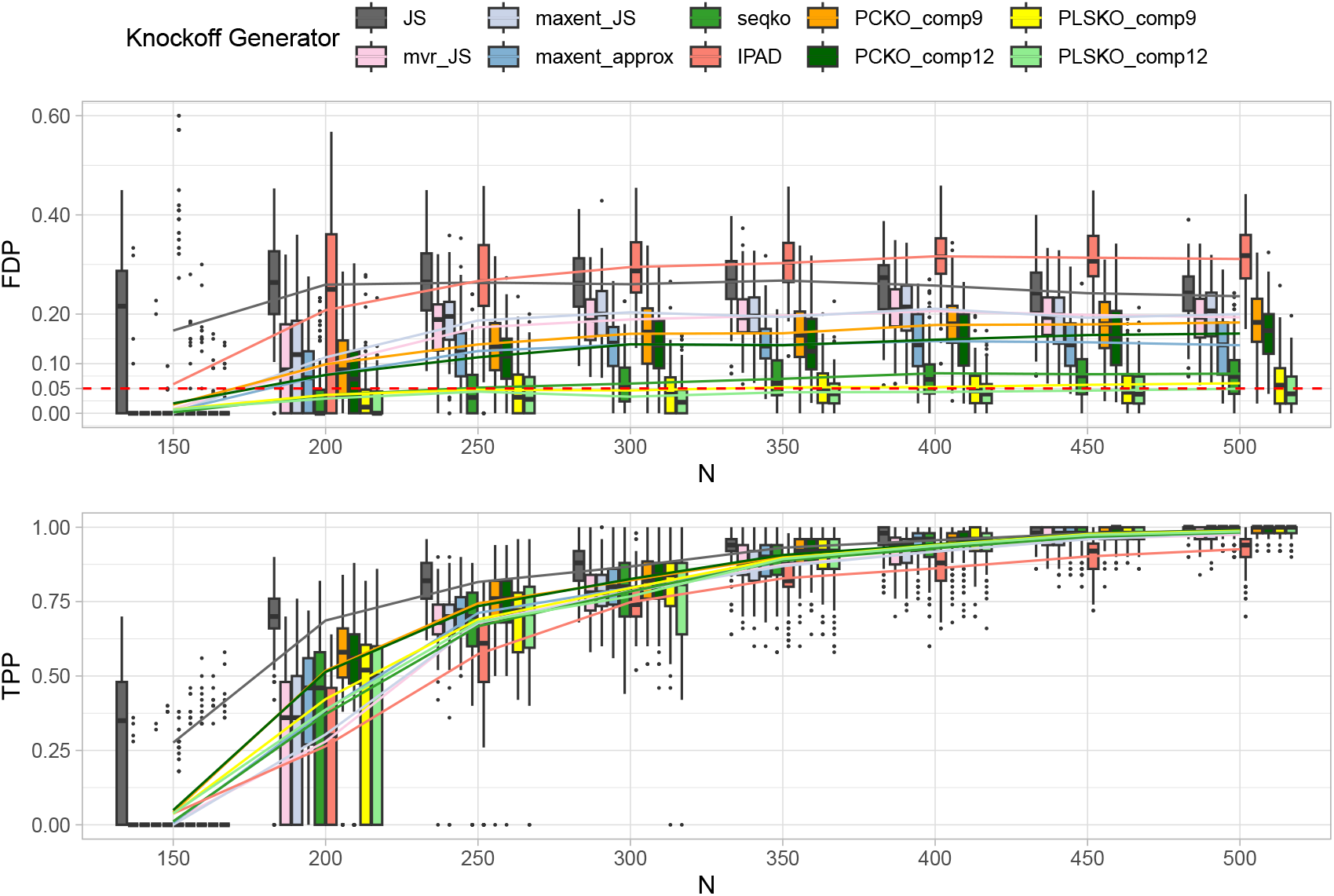
Simulations: FDR and power of PLSKO and other knockoff generators when **X** is generated from a quadratic factor model w.r.t sample size, with *p* = 500, 5 blocks, 3 latent factors, 10% signal proportion and a target FDR of 0.05 across 100 replications. FDP: false discovery proportion; TPP: true positive proportion. Solid lines represent the mean of FDP and TPP, i.e., the estimates FDR and power.

## 4 Semi-Simulation Studies

Assuming that biological data follow a Gaussian or factor model might not be realistic. We showed in our simulation results that violation of assumption from knockoff generators, or poor fitness can lead to inflated FDR. Thus, we next used real biological data **X**, but we simulated *y*, and checked the validity and robustness of knockoff generators, including PLSKO, on different types biological data with unknown distribution.

### Semi-simulation setting

We performed semi-simulations based on the two case-control studies of preeclampsia. After data preprocessing, we ran the semi-simulation for each of the datasets over 50 repetitions. In each run, we first randomly selected a subset of variables **X**^**subset**^ to avoid zero power in high-dimensional data and to retain the joint distribution of the original **X**. Then, similar to our simulations, we simulated *y* from a set of randomly selected important variables from **X**^**subset**^. Once the knockoff variables for **X**^**subset**^ were generated, we applied knockoff filtering with a target level of FDR (cfRNA:0.05, multi-omic datasets:0.10); and used FDP and TPP as a measure of performances from each repetition.

### Simulation results

For datasets from both cfRNA and multi-omics studies, we found that both PLSKO-full and PLSKO-full-sparse were the only methods controlling for the modified FDR (mFDR) close to the controlled level, regardless of the number of variables in **X**^**subset**^ (cfRNA: Figure 3, multi-omics: Supplemental Figures S6b, S7a and S7b). The other methods, including PLSKO with arbitrary and liberal parameters (3 components and 80% top correlation) had inflated mFDR around 0.2. We determined the number of components in PLSKO-full and PLSKO-full-sparse based on the *PC*_*p*1_ criterion from Bai and Ng (2002), ranging from 4 to 8 with a mode of 6 in the repeatedly sampled **X**^**subset**^. Despite the inherent trade-off between FDR and power, both methods were able to generate knockoff variables with the lowest FDR and modestly enhanced power. Additionally, PLSKO-full-sparse performed similarly to PLSKO-full but only included 20% of variables on each component for fitting the conditional distribution, suggesting that sparse PLS regression did not cause under-fitting. Since RNA-seq counts are assumed to marginally follow a negative binomial distribution (Robinson and Smyth, 2008), which violates the assumption in SOA methods, we further normalised the RNA-seq data to ensure that **X**^**subset**^ follow a Gaussian distribution. However, we still found that the FDR was inflated and power was low (Supplemental Figure S6a), suggesting poor fitness of covariance and information loss.

**Figure 3:**
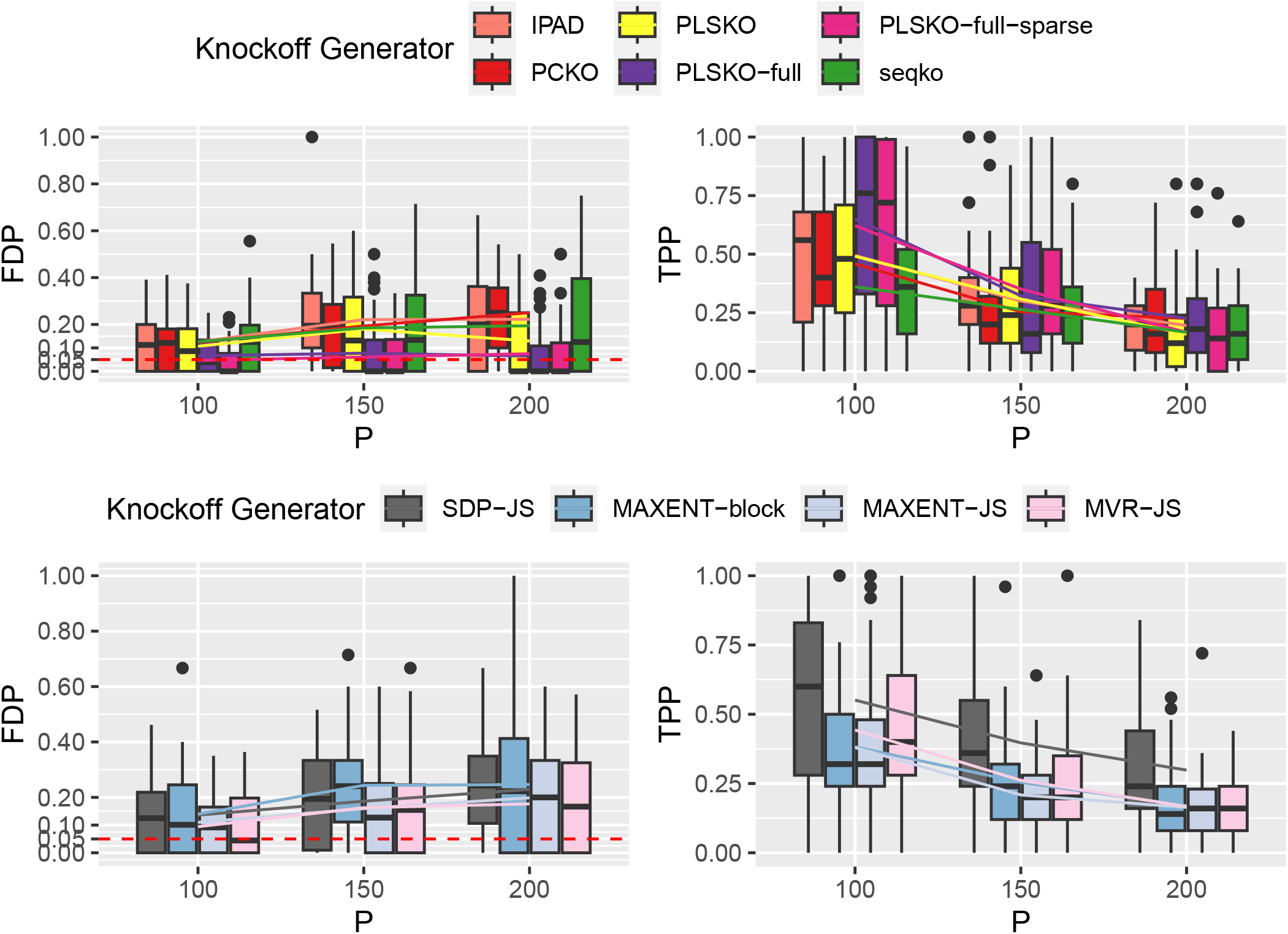
Semi-simulations: FDR and power of PLSKO and other knockoff generators on cell-free RNA-seq data w.r.t numbers of variables, with target FDR = 0.05, sample size *n* = 71, number of important variables *p*_*s*_ = 25, over 50 repetitions. Upper panel: SCIP-based methods and IPAD; lower panel: SOA methods. FDP: false discovery proportion; TPP: true positive proportion. Solid lines represent mean of FDP and TPP, i.e., the estimated FDR and power.

In summary, both PLSKO-full and PLSKO-full-sparse with no prior neighbour screening and a number of components determined by the *PC*_*p*1_ criterion were robust to different types of real biological data without over-simplifying the assumptions on **X**. We found that the other knockoff generators and PLSKO with arbitrary liberal setting failed due to insufficient fitness on these complex data.

## 5 Case Studies

We then applied PLSKO on the preeclampsia cfRNA and multi-omics studies, where the response variable is preeclampsia or not. We first reduced the number of variables in the datasets by focusing solely on circulating placenta-elevated genes and proteins. As abnormal placenta is central to preeclampsia, our hypothesis is that this subset of genes and proteins might reflect changes from placenta. In the cfRNA dataset, we selected 81 placenta-elevated genes defined according to The Human Protein Atlas (Uhlén et al., 2015). In the proteomics dataset, we kept 36 placenta-released proteins from plasma according to Degnes et al. (2022). For the microbiome data, we kept 144 Operational Taxonomic Units (OTUs) filtered with a sum of counts > 0.01% of the total sum of counts.

We generated knockoff variable sets for each dataset using PLSKO-full with no neighbour filtering and 8 PLS components. The LCD generated from logistic lasso regression was used as importance statistic. To accommodate for the randomness of knockoff generators, we ran PLSKO-full 50 times and incorporated the knockoff variable sets using Aggregation knockoff (AKO) with target FDR level 0.05. Although generating multiple knockoffs is not necessary, it is highly recommended as AKO improves stability and power, while the selection frequency from the repeated single-run knockoffs from PLSKO can be used as a measure of selection stability. We compared our selection with two commonly-used methods for differential expression (DE), *limma* (Smyth, 2005) and Wilcoxon test with BH procedure (Table 1).

**Table 1:** Genes, proteins from plasma and OTUs (reported either at the species ‘s_’, family ‘f_’ or genus ‘g_’ levels) identified as frequently selected by PLSKO-full run 50 times (frequency > 0.1), and results aggregated with PLS-AKO. Selections are compared with differential expression (DE) tests limma and Wilcoxon.

| Dataset | Predictor | Selected Feature | PLSKO frequency | PLS-AKO selection | Declared DE |
| --- | --- | --- | --- | --- | --- |
| cell-free RNA-seq (n = 71) | Placenta-specific elevated gene expression (p = 81) | PHACTR2 | 0.92 | Yes | No |
|  |  | TENT5A/<br>FAM46A | 0.88 | Yes | Yes (limma, Wilcoxon) |
|  |  | GSE1 | 0.82 | Yes | No |
|  |  | MBNL3 | 0.58 | Yes | Yes (Wilcoxon) |
|  |  | HEMGN | 0.56 | Yes | No |
|  |  | MAFK | 0.46 | No | No |
|  |  | BPGM | 0.36 | No | Yes (Wilcoxon) |
|  |  | AFF1 | 0.3 | No | No |
|  |  | CSF2RB | 0.24 | No | No |
|  |  | TRIM10 | 0.14 | No | No |
|  |  | COBLL1 | 0.1 | No | No |
| Proteomics (n = 36) | Placenta-released proteins (p = 36) | SERPINE1 | 0.32 | Yes | No |
|  |  | HSPB1 | 0.3 | Yes | No |
|  |  | DKK1 | 0.26 | No | No |
|  |  | CXCL10 | 0.1 | No | No |
| Microbiome study (n = 49) | 144 OTUs | g_ Corynebacterium | 0.26 | Yes | No |
|  |  | s_ F. nucleatum | 0.26 | Yes | No |
|  |  | f_ Neisseriaceae | 0.2 | No | No |
|  |  | s_ L. acidophilus | 0.1 | No | No |

In the cfRNA data, 11 genes were selected more than 10% of the time across the 50 repetitions by PLSKO. Amongst those genes, 5 were selected by PLS-AKO, whereas *limma* selected 1 and the Wilcoxon test 3 of these genes. The most frequently gene selected by PLSKO and PLS-AKO was PHACTR2, which not declared significantly associated with preeclampsia by either limma or a Wilcoxon test. However, when we controlled for the second most frequently selected gene TENT5A using a logistic regression, PHACTR2 was declared significant (Wald test). The BPGM gene was declared DE by Wilcoxon, but was not identified by PLS-AKO and selected by PLSKO relatively infrequently. This could be due to the high collinearity of MBNL3 (fourth most frequent) and BPGM, as the latter was not found associated with preeclampsia when controlling on MBNL3.

These results highlight the difference of controlled variable selection (knockoff filtering) and marginal testing (DE analysis): variables selected via controlled variable selection are not necessarily DE; and controlled variable might not be selected with collinear variables in some cases.

In each of the multi-omic studies, PLSKO and PLS-AKO selected several biological variables missed by the DE tests. All proteins selected by PLSKO have been reported to be related to preeclampsia, including SERPINE1 (Zhao et al., 2013), HSPB1 (Martin et al., 2022), CXCL10 (Gotsch et al., 2007), DKK1 (Kasoha et al., 2021) and so were the selected OTUs including, the species *Fusobacterium Nucleatum* (Barak et al., 2007) and the family *Neisseriaceae* (Taylor et al., 2023). Results with various types of filtering or the whole datasets are presented in Supplemental Tables S1 and S2.

Overall, our application of PLSKO in biological data sets showed that our method is able to identify biologically relevant omics features with higher power than marginal DE_tests_.

## 6 Discussion

PLSKO is a new knockoff variable generator for biological high-throughput data. It is a filtering approach leading to the selection of important variables related to a response of interest with FDR control. PLSKO can be paired with various variable selection models depending on the type of analysis and biological questions. Our results in simulations, semi-simulations and applications in multiple types of omics data (RNA-seq, proteomics, microbiome) show that PLSKO is more robust in FDR control and has higher statistical power than existing knockoff generators. Compared to classic DE analysis, we also showed that PLSKO selected more variables that were biologically highly relevant.

In the case studies we presented, we filtered subsets of variables before applying PLSKO due to their limited sample size. Future extensions of PLSKO could include group-PLSKO, which relaxes control within variable groups, addressing issues of low power in extreme high-dimensional settings. In addition, since PLS is amenable to omics data integration, PLSKO could also be extended for the integration of multi-omics studies to enhance insights and reproducibility in complex diseases.

## Availability of data and codes

All analyses were conducted using the `R` software and are fully reproducible using the codes available on Github https://github.com/esheeep/PLSKO. The `R` packages used in our analyses are listed in Supplementary Material S3.9.

## Author contributions statement

GY and KALC designed the study and wrote the manuscript. GY developed the method, conducted the analysis and interpreted the results, under the supervision of KALC, EM and ED. EM and ED provided feedback on the case studies. All authors read and approved the final manuscript.

