## Supplemental for "PLSKO: a robust knockoff generator to control false discovery rate in omics variable selection"

### S1 Supplementary Figures

#### S1.1 Method diagram

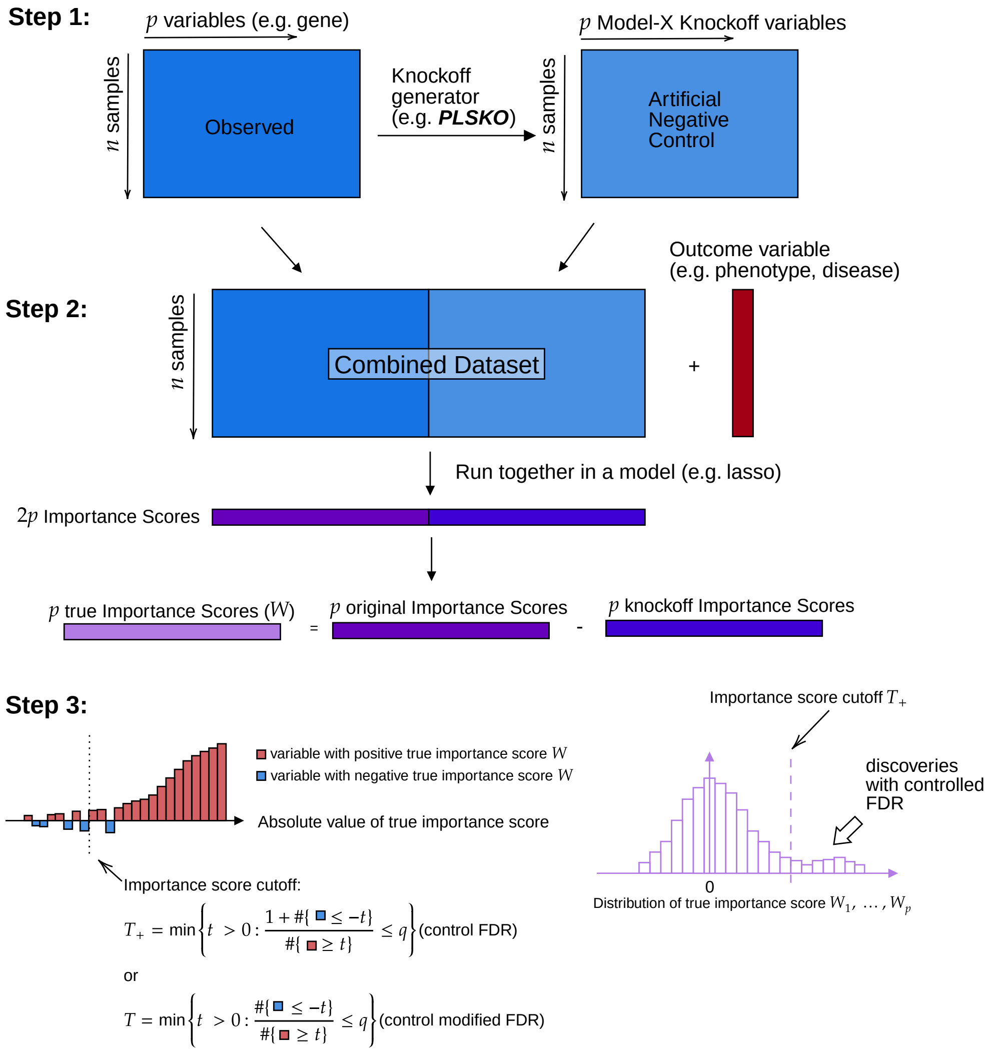


Figure S1: Overview of knockoff filtering. Step 1: Knockoff variable construction: we generate knockoff variables that follow model-X properties as the artificial negative control. Step 2: we run the original and knockoff variables together to calculate importance score that measure the *true* importance of each variable to the response variable $y$ by contrasting it with its knockoff. For example, the lasso coefficient difference (LCD) can be used, but other models and importance statistics that follow the *flip-sign* property are also applicable. Step 3: we find the threshold controlling the large negative importance score proportion that is less than the nominal FDR level and select the variables with larger importance scores.

#### S1.2 Simulation results

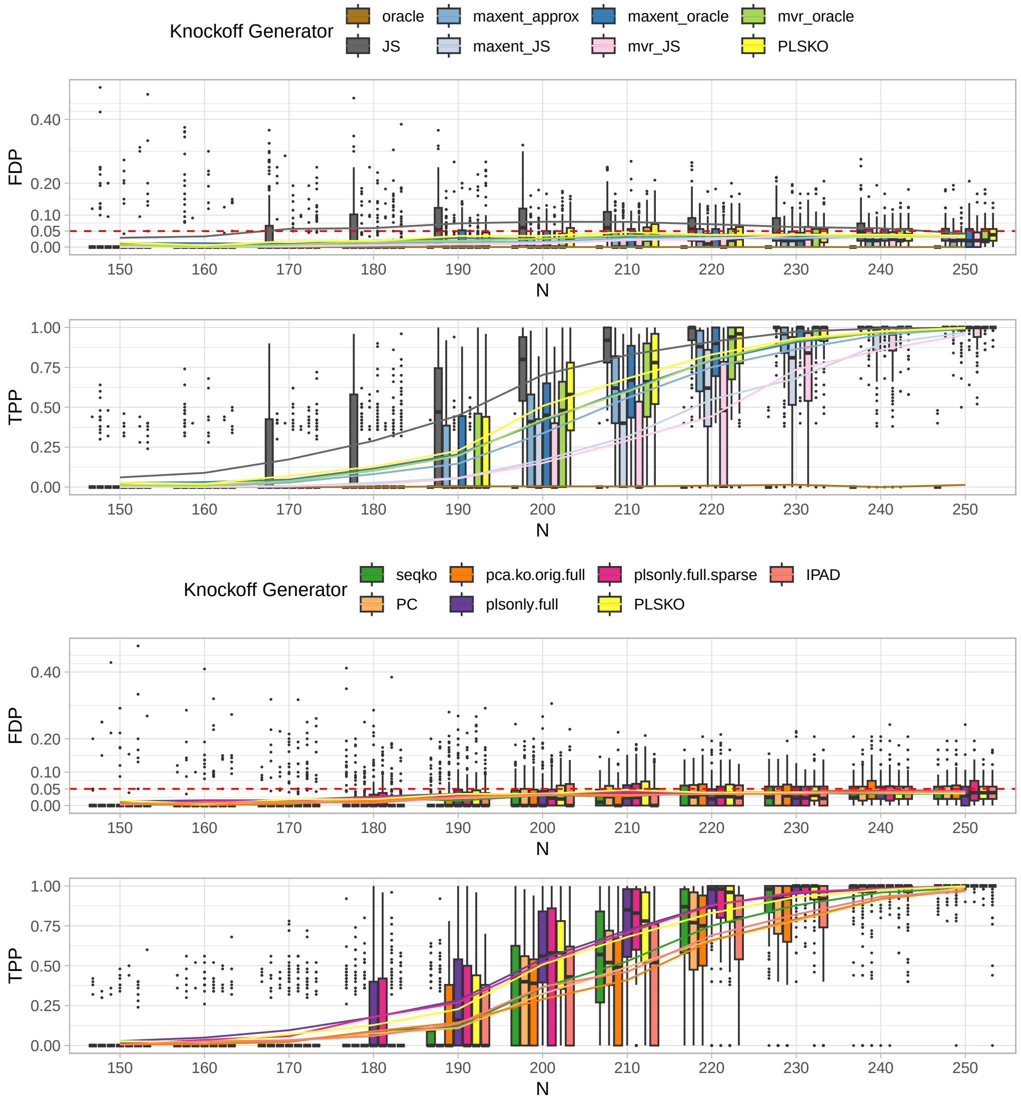


Figure S2: Simulation results: FDR and power of PLSKO and other knockoff generators when **X** follows a Gaussian factor model w.r.t sample size, with $p = 500$, 5 blocks, 3 latent factors, 10% signal proportion and a target FDR of 0.05 across 100 replications. FDP: false discovery proportion; TPP: true positive proportion. Solid lines represent the mean of FDP and TPP, i.e., the estimates FDR and power.
Upper panel: PLSKO and second-order approximation (SOA) methods; Lower panel: PLSKO and other SCIP-based methods. Oracle: SDP with the true covariance; JS: SDP with the James-Stein type shrunk covariance; PC: SCIP with PC regression and neighbour presceening; pca.ko.orig.full: SCIP with PC regression without neighbour prescreening; plsonly.full: PLSKO-full with PLS regression without neighbour screening; plsonly.full.sparse: PLSKO-full-sparse without neighbour screening and sparse PLS regression with 100 variables (20% of $p$) kept on each component. All PC or PLS regression components are equal to 3.
The SOA-SDP method, which is an exact knockoff generator with perfect 'oracle' knowledge of the covariance, had near-zero power. This phenomenon aligns with the findings from Spector and Janson (2022) that SDP method loses power when the rank of the gram matrix ($G = X^\top X$) is less than $2p$, and the rank is $2r << 2p$ in this factor model simulation. Most of these methods generated valid knockoff variables that controlled FDR when $X$ follows the Gaussian factor model, while PLSKO and its variants showed a slight advantage in power.

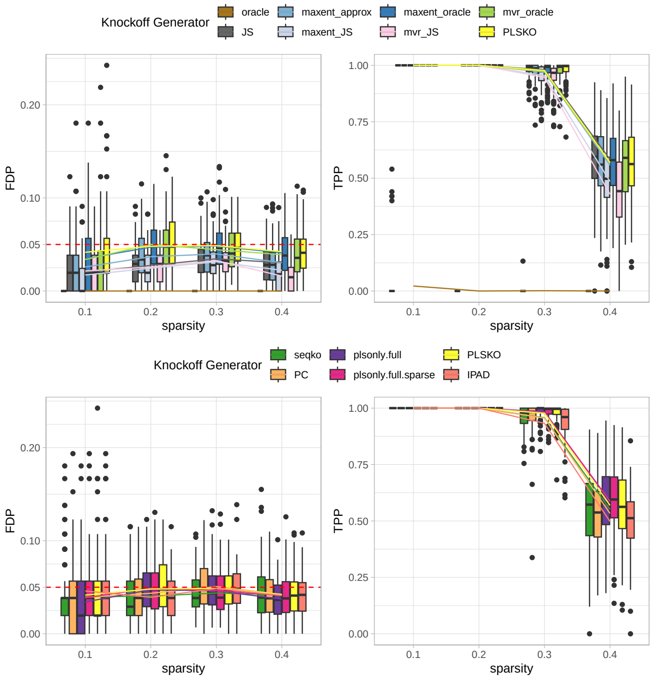


(a) Simulation with varying proportions of important variables.

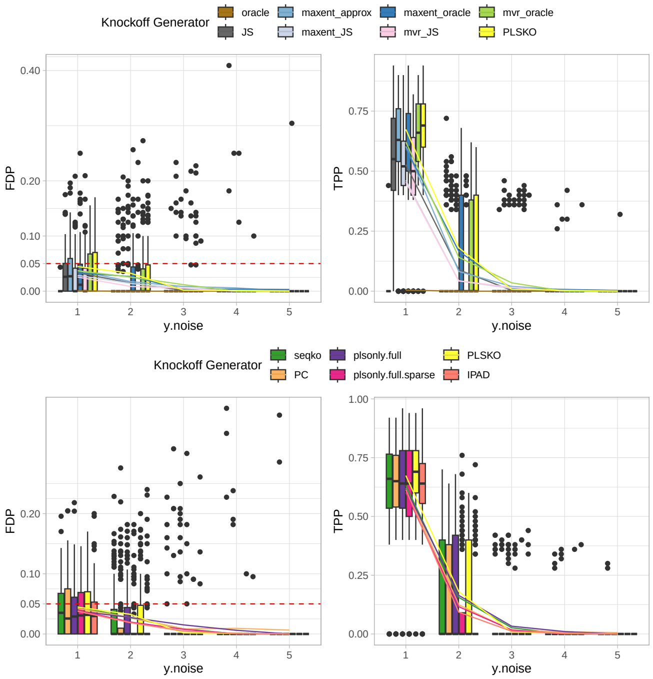


(b) Simulation with varying ratio of signal and noise (measured and unmeasured important variables)

Figure S3: Simulation on block factor model with $y$ generated when $\mathbf{X}$ follows a Gaussian factor model w.r.t (a) varying proportions of important variables $p_s$ and (b) varying noise added to $y$, with $n = 500, p = 500$, 5 blocks, 3 latent factors, 10% signal proportion and a target FDR of 0.05 across 100 replications. Upper panel: PLSKO and SOA methods; Lower panel: PLSKO and other SCIP-based methods. Sparsity represents the proportion of important variables, that is, 0.1 represents 10% of $p$ variables used to generate $y$, and so on. y.noise ($c$): a random noise $\epsilon \sim \mathcal{N}(0, p_s \times c^2)$ is added to $y$, representing the ratio of noise to signal, that is when $c = 1$ the noise signal ratio is 16:1.

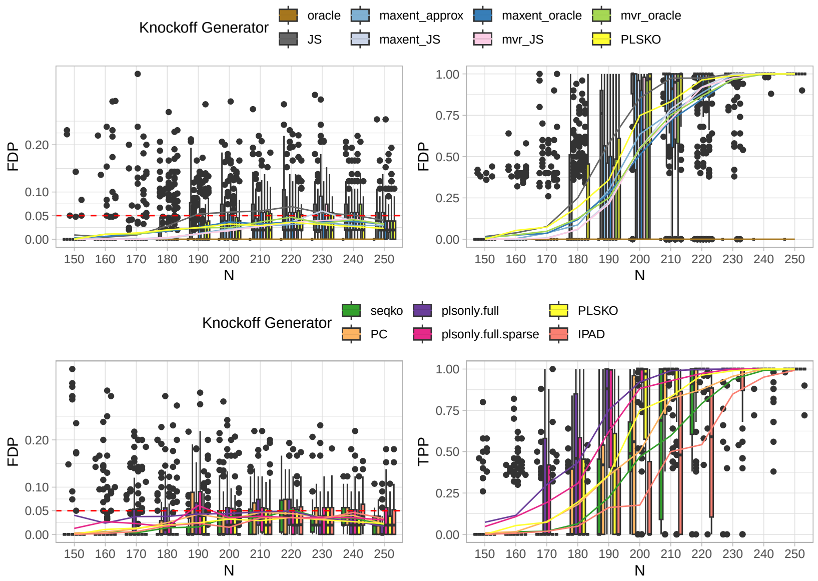


(a) Simulation with varying sample size in multivariate Gaussian distribution with block equi-correlated covariance ($\rho = 0.5$)

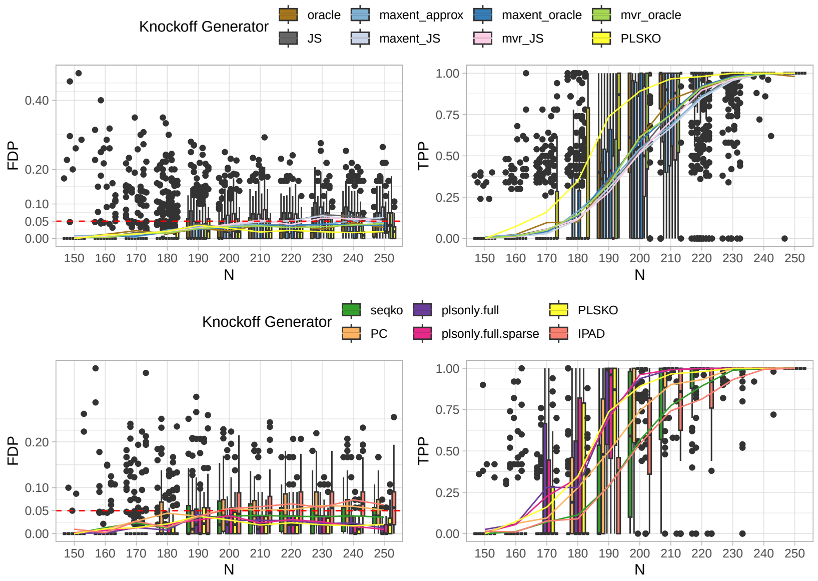


(b) Simulation with varying sample size in multivariate Gaussian distribution with block AR1 covariance ($\rho = 0.5$)

Figure S4: Simulation experiments on multivariate Gaussian distribution with (a) block equi-correlated covariance and (b) AR1 covariance, across over varying sample sizes, with $p = 500$, 5 blocks, 3 latent factors, 10% signal proportion and a target FDR of 0.05 across 100 replications.

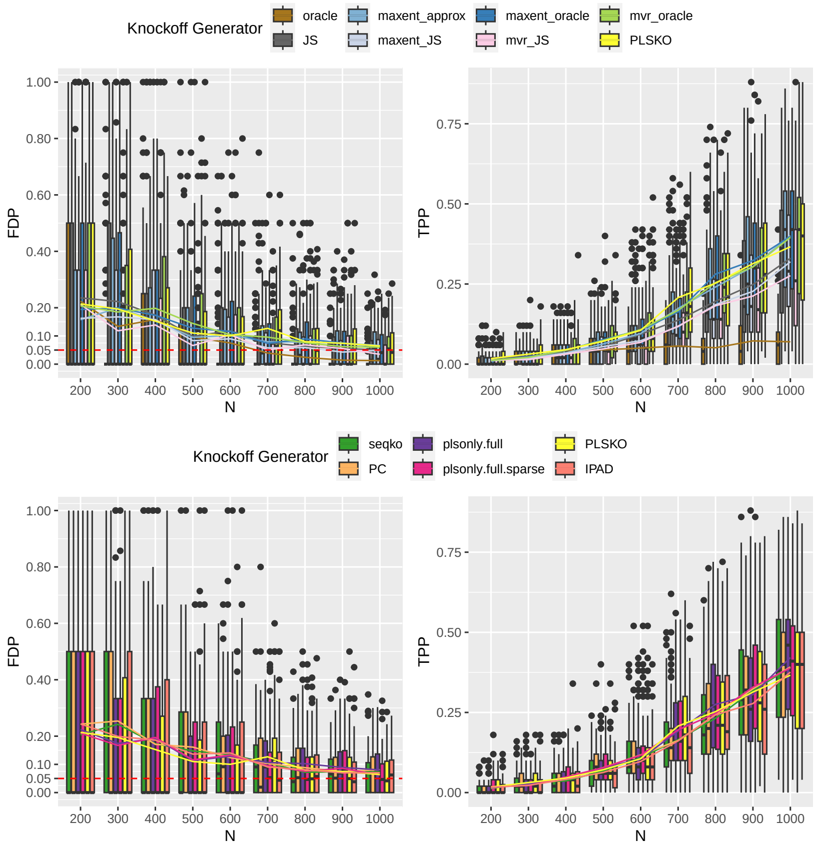


Figure S5: Simulation with varying sample size with categorical $y$ (controlled on modified FDR) when $X$ follows Gaussian factor model with with $p = 500$, 5 blocks, 3 latent factors, 10% signal proportion, $y$ generated by Sigmoid function of the linear combination of signal variables and a target FDR of 0.05 across 100 replications. Lasso logistic regression and the coefficient difference are used as the importance score. Lower power showed than LCD on linear generated $y$ as categorical $y$ contains fewer information. Modified FDR are used and controlled here since FDR controlled by the conservative threshold $T_+$ leads to no power. PLSKO and its variants perform similar with other knockoff generators.

#### S1.3 Semi-simulations results

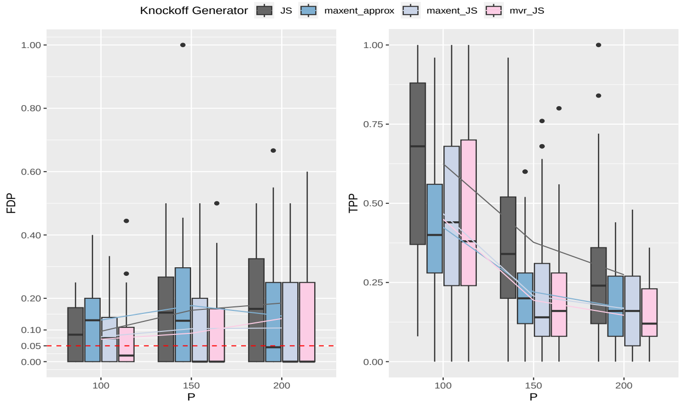


(a) Semi-simulation of second-order approximation knockoff generators on marginally-normalised cfRNA data across varying numbers of variables, with target (modified) FDR = 0.05, sample size $n = 71$, number of important variable $p_s = 25$ over 50 repetitions.

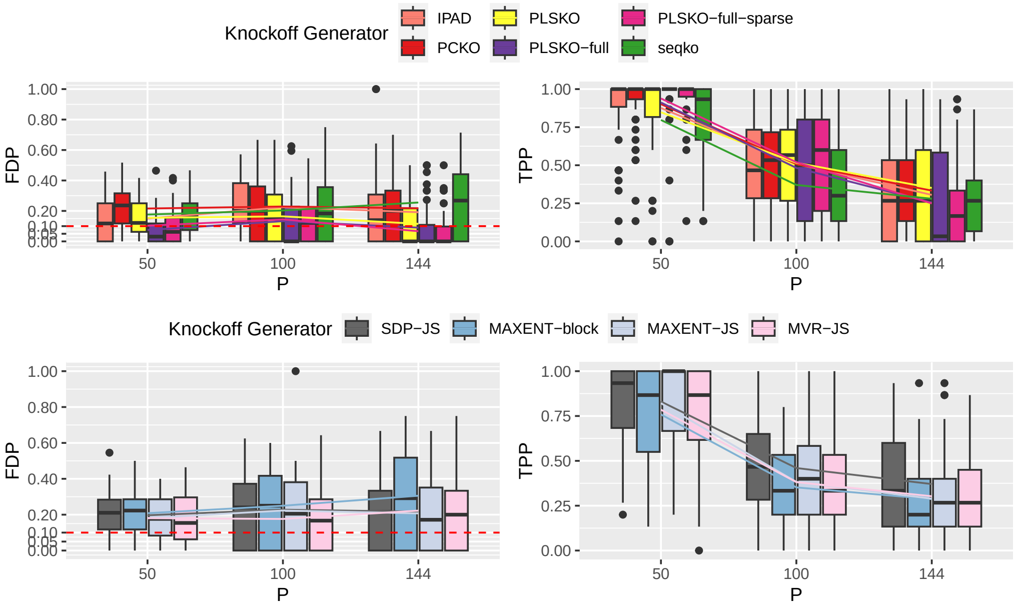


(b) Semi-simulations: FDR and power of PLSKO and other knockoff generators on microbiome data across varying numbers of variables, with target FDR = 0.10, sample size $n = 49$, number of important variable $p_s = 15$ over 50 repetitions. Upper panel: SCIP-based methods and IPAD; lower panel: SOA methods. FDP: false discovery proportion; TPP: true positive proportion. Solid lines represent mean of FDP and TPP, i.e., the estimates FDR and power. Modified FDR are used since FDR controlled by the conservative threshold $T_+$ leads to no power.

Figure S6

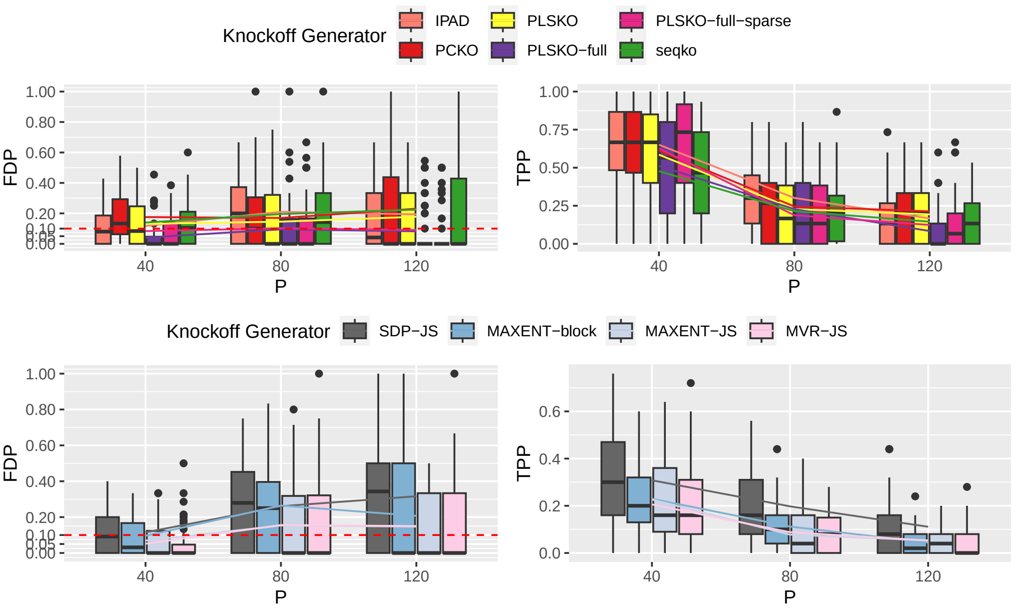


(a) Semi-simulations: FDR and power of PLSKO and other knockoff generators on proteome data across varying numbers of variables, with target FDR = 0.10, sample size $n = 36$, number of important variable $p_s = 15$ over 50 repetitions. Upper panel: SCIP-based methods and IPAD; lower panel: SOA methods. Modified FDR are used since FDR controlled by the conservative threshold $T_+$ leads to no power.

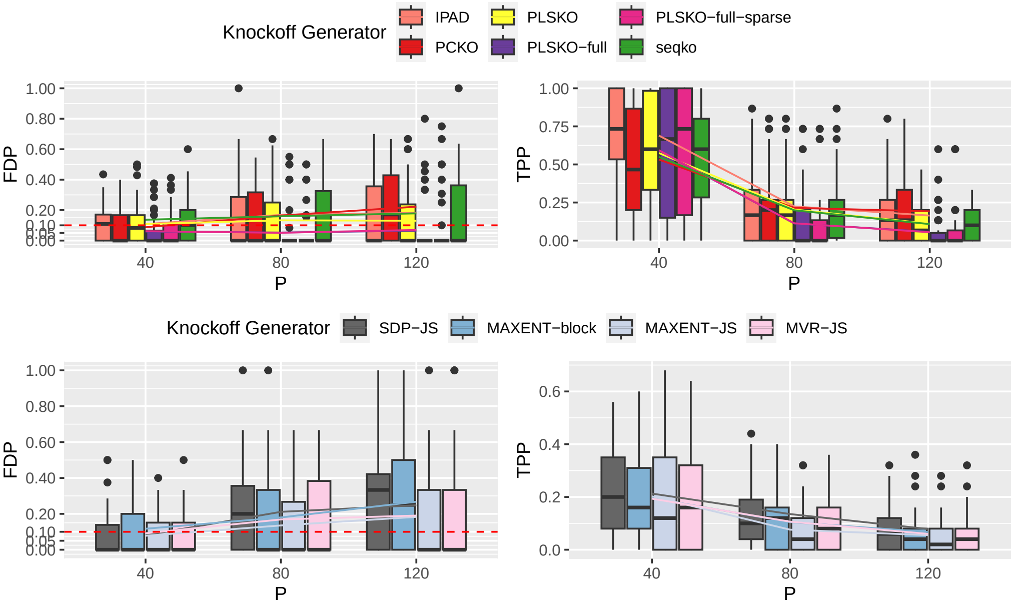


(b) Semi-simulations: FDR and power of PLSKO and other knockoff generators on the urine metabolome data across varying numbers of variables, with target FDR = 0.10, sample size $n = 36$, number of important variable $p_s = 15$ over 50 repetitions. Upper panel: SCIP-based methods and IPAD; lower panel: SOA methods. FDP: false discovery proportion; TPP: true positive proportion. Solid lines represent mean of FDP and TPP, i.e., the estimates FDR and power.

Figure S7

#### S2 Supplementary Tables

Table S1: Selected preeclampsia-related features from alternative subsets of the omics data over 50 generated knockoffs

| **Dataset** | **Predictors** | **Selected Feature** | **PLSKO frequency** | **Selected by PLS-AKO** | **DE by other methods** |
|---|---|---|---|---|---|
| cell-free Transcriptomics from multi-omics data (n = 36) | Placenta-specific elevated gene expression (p = 77) | IGF2 | 0.04 | Yes | No |
| | | BPGM | 0.04 | No | No |
| | | HBM | 0 | No | Yes (limma) |
| | Placenta-specific (fold-change > 3) elevated gene (p = 86) (HPA v23) | IGF2 | 0.06 | Yes | No |
| | | EMILIN2 | 0.06 | No | No |
| cell-free Transcriptomics (n = 71) | Placenta-specific elevated gene expression (p = 96) (HPA v23) | MBNL3 | 0.44 | Yes | Yes (Wilcoxon) |
| | | CD36 | 0.42 | Yes | No |
| | | PHACTR2 | 0.18 | No | No |
| | | FLI1 | 0.12 | No | No |
| | | CDK6 | 0.1 | No | No |
| | Genes with the highest variance (p = 200) | CA1 | 0.42 | Yes | No |
| | | RN7SL564P | 0.38 | Yes | Yes (limma, Wilcoxon) |
| | | RN7SL381P | 0.34 | Yes | Yes (limma, Wilcoxon) |
| | | MAGI2-AS3 | 0.3 | No | No |
| | | RN7SL736P | 0.14 | No | Yes (limma, Wilcoxon) |
| | | RNA5SP387 | 0.1 | No | No |
| Proteomics from multi-omics study (n = 36) | Placenta-specific elevated gene expression (p = 101) (HPA v23) | IL1RAP | 0.36 | Yes | No |
| | | COLEC12 | 0.36 | Yes | No |
| | | HAPLN1 | 0 | No | Yes (Wilcoxon) |
| | Placenta-specific elevated gene expression (p = 63) (HPA v19) | IL1RAP | 0.58 | Yes | Yes (limma) |
| | | LEP | 0.58 | Yes | Yes (limma, Wilcoxon) |
| | | GSTA3 | 0.1 | No | No |
| | | HAPLN1 | 0 | No | Yes (Wilcoxon) |
| Microbiome data from a multi-omics study (N = 49) | 144 OTUs (TMM normalisation) | s_L. acidophilus | 0.04 | Yes | No |
| | | s_L. iners | 0.04 | No | No |

Table S2: Selected variables in the cell-free RNA dataset (without subselection on placenta elevated variables) by PLSKO, limma and Wilcoxon test [1]

| **Datasets: cell-free RNA , 3,000 genes with the highest variance, n = 71** | | | | | |
|---|---|---|---|---|---|
| **limma significant** | | **PLSKO** | | **Frequency** | **PLSKO-AKO** |
| ENSG00000211896 | IGHG1 | ENSG00000263968 | RN7SL381P | 0.22 | T |
| ENSG00000239437 | RN7SL752P | ENSG00000240606 | RN7SL564P | 0.18 | F |
| ENSG00000239607 | RN7SL573P | ENSG00000119048 | UBE2B | 0.08 | F |
| ENSG00000239899 | RN7SL674P | | | | |
| ENSG00000240606 | RN7SL564P | | | | |
| ENSG00000240869 | RN7SL128P | | | | |
| ENSG00000241529 | RN7SL767P | | | | |
| ENSG00000243352 | RN7SL8P | | | | |
| ENSG00000244642 | RN7SL396P | | | | |
| ENSG00000251705 | RNA5-8SP6 | | | | |
| ENSG00000263968 | RN7SL381P | | | | |
| ENSG00000264169 | RN7SL665P | | | | |
| ENSG00000264275 | RN7SL753P | | | | |
| ENSG00000264916 | RN7SL230P | | | | |
| ENSG00000264978 | RN7SL630P | | | | |
| ENSG00000265735 | RN7SL5P | | | | |
| ENSG00000266439 | RN7SL493P | | | | |
| ENSG00000266794 | RN7SL7P | | | | |
| ENSG00000275803 | RN7SL736P | | | | |

| **Datasets: cell-free RNA, all 7,160 genes after prefiltering, n = 71** | | | | | |
|---|---|---|---|---|---|
| **limma** | | **PLSKO** | | **Frequency** | **PLSKO-AKO** |
| ENSG00000013561 | RNF14 | ENSG00000263968 | RN7SL381P | 0.2 | F |
| ENSG00000239437 | RN7SL752P | ENSG00000013561 | RNF14 | 0.18 | F |
| ENSG00000239607 | RN7SL573P | ENSG00000103342 | GSPT1 | 0.18 | F |
| ENSG00000239899 | RN7SL674P | ENSG00000240606 | RN7SL564P | 0.16 | F |
| ENSG00000240606 | RN7SL564P | ENSG00000109787 | KLF3 | 0.12 | F |
| ENSG00000240869 | RN7SL128P | ENSG00000163875 | MEAF6 | 0.12 | F |
| ENSG00000241529 | RN7SL767P | | | | |
| ENSG00000243352 | RN7SL8P | | | | |
| ENSG00000244642 | RN7SL396P | | | | |
| ENSG00000263968 | RN7SL381P | | | | |
| ENSG00000264169 | RN7SL665P | | | | |
| ENSG00000264275 | RN7SL753P | | | | |
| ENSG00000264916 | RN7SL230P | | | | |
| ENSG00000264978 | RN7SL630P | | | | |
| ENSG00000265735 | RN7SL5P | | | | |
| ENSG00000266439 | RN7SL493P | | | | |
| ENSG00000275803 | RN7SL736P | | | | |

| **Datasets: cell-free RNA, 200 variables with the highest variance, n = 71** | | | | | | | |
|---|---|---|---|---|---|---|---|
| **limma** | | **PLSKO** | | **Frequency** | **PLSKO-AKO** | **Wilcoxon** | |
| ENSG00000200434 | RNA5-8SP2 | ENSG00000133742 | CA1 | 0.42 | T | ENSG00000239437 | RN7SL752P |
| ENSG00000239437 | RN7SL752P | ENSG00000240606 | RN7SL564P | 0.38 | T | ENSG00000239607 | RN7SL573P |
| ENSG00000239607 | RN7SL573P | ENSG00000263968 | RN7SL381P | 0.34 | T | ENSG00000239899 | RN7SL674P |
| ENSG00000239899 | RN7SL674P | ENSG00000234456 | MAGI2-AS3 | 0.3 | F | ENSG00000240606 | RN7SL564P |
| ENSG00000240606 | RN7SL564P | ENSG00000275803 | RN7SL736P | 0.14 | F | ENSG00000240869 | RN7SL128P |
| ENSG00000240869 | RN7SL128P | ENSG00000201096 | RNA5SP387 | 0.1 | F | ENSG00000241529 | RN7SL767P |
| ENSG00000241529 | RN7SL767P | | | | | ENSG00000243352 | RN7SL8P |
| ENSG00000243352 | RN7SL8P | | | | | ENSG00000244230 | RN7SL151P |
| ENSG00000244230 | RN7SL151P | | | | | ENSG00000244642 | RN7SL396P |
| ENSG00000244642 | RN7SL396P | | | | | ENSG00000251705 | RNA5-8SP6 |
| ENSG00000251705 | RNA5-8SP6 | | | | | ENSG00000263968 | RN7SL381P |
| ENSG00000263968 | RN7SL381P | | | | | ENSG00000264169 | RN7SL665P |
| ENSG00000264169 | RN7SL665P | | | | | ENSG00000264275 | RN7SL753P |
| ENSG00000264275 | RN7SL753P | | | | | ENSG00000264916 | RN7SL230P |
| ENSG00000264916 | RN7SL230P | | | | | ENSG00000264978 | RN7SL630P |
| ENSG00000264978 | RN7SL630P | | | | | ENSG00000265735 | RN7SL5P |
| ENSG00000265735 | RN7SL5P | | | | | ENSG00000266439 | RN7SL493P |
| ENSG00000266439 | RN7SL493P | | | | | ENSG00000275803 | RN7SL736P |
| ENSG00000275803 | RN7SL736P | | | | | | |

[1] a. No DEG were detected by Wilcoxon test in either 3,000 genes or the whole dataset; b. Pseudogenes (e.g. RN7SL752P) tended to be selected together by the marginal tests as the genes are highly correlated (e.g. correlation > 0.85), while PLSKO selected only one or two pseudogenes as other pseudogenes have been controlled;

### S3 Supplementary Methods

#### S3.1 Overview of Model-X knockoff filter

Barber and Candès (2015) introduced a novel framework, "knockoffs," for obtaining FDR control in feature selection, bypassing the standard calculation of p-values. The knockoff filtering procedure can be used as a wrapper by combining it with any feature selection method that generates feature importance measures that meet certain conditions. The basic principle of the knockoff procedure is to create a (or multiple) artificial "knockoff" data based on the original data, which is similar to the original real data, without using any information from the response variable. Then, the real data ($X$) and their knockoff copy data ($\widetilde{X}$) are run together into a model to obtain an importance measure for each original or knockoff variable so that the knockoff variables then function as negative controls for the original covariates. Each original variable's estimated importance measure is then compared to that of its corresponding knockoff variable, whose actual effect is known to be zero. Variables are chosen on the basis of this means of variable importance, which is low when the influence of the real variable cannot be distinguished from the estimated effect of a knockoff variable.

Knockoff filtering aims to select 'relevant' variables that are conditionally dependent on the response variable given all the other covariates, i.e., a subset of $X$ that affects $Y$. $X_j$ is referred to as 'null' if it is conditionally independent of the response variable $Y$ once the other $p-1$ variables are given.

Knockoff filtering for the selection of important variables consists of three main steps:

**Step 1: Knockoff variable construction.** When $n \leq p$, the knockoff variables $\widetilde{X}$ can be generated as Model-X knockoffs, which are defined as a new set of random variable $\widetilde{X_j}, j = 1, \cdots, p$ that satisfies the properties:
1) for any subset $S \subset \{1, \cdots, p\}$,

$$(X, \widetilde{X})_{\text{swap}(S)} \stackrel{d}{=} (X, \widetilde{X}), \tag{1}$$

where $(X, \widetilde{X})_{\text{swap}(S)}$ is obtained by swapping the $X_j$ and $\widetilde{X}_j$ for each $j \in S$;
2) $\widetilde{X} \perp\!\!\!\perp Y|X$ (guaranteed if $\widetilde{X}$ is constructed without looking at $Y$).

Existing approaches to construct valid model-X knockoff will be discussed in the next section S3.2.

**Step 2: Calculate the important statistics.** Once the knockoff variables are generated, the model for variable selection is run on an augmented data set with response $y$ and $2p$ many features $X_1, ..., X_p, \widetilde{X}_1, ..., \widetilde{X}_p$. Then, a pairwise statistics $W_j$ for each $j \in \{1, ..., p\}$ is computed. This statistic depends on the response, original variables, and knockoffs. A valid statistic should have *flip-sign property* that swapping the $j$th variable with its knockoff has the effect of changing the sign of $W_j$ (Candès et al., 2018). A large positive value of $W_j$ provides some evidence that the distribution of Y depends upon $X_j$, whereas under the null (for those null/noise variables), $W_j$ has a symmetric distribution and, therefore, is equally likely to take on positive and negative values. For example, a common option for valid $W_j$ can be the lasso coefficient difference (LCD) (with the value of $\lambda$ can be decided by cross-validation).

**Step 3: Find the threshold and select variables.** Based on the property of pairwise exchangeability (see Candès et al. (2018) Section 3.2) where the $W_j$ for noise variables are equally likely to be positive or negative (distributed symmetrically to zero, i.e. $\#\{null\ j : W_j \leq -t\} \stackrel{d}{=} \#\{null\ j : W_j \geq t\}$), the set of important features with an FDR at a pre-specified level $q \in [0, 1]$ is selected as $\hat{S} = \{j : W_j \geq t\}$ with the threshold $t = T$, defined as:

Table S3: Selected genes co-expressed with gene RNF14 from the placenta-elevated genes in the cell-free plasma data. The gene RNF14 was identified as DE in preeclampsia cell-free plasma (Table S2). Marginal DE tests are not applicable to this case with a non-categorical response variable.

| Dataset | Predictors | Selected Feature | Frequency by PLSKO | Selected by PLS-AKO |
|---|---|---|---|---|
| Cell-free transcriptomics (n = 71) | Placenta-specific elevated gene (p = 81) | BPGM | 0.84 | Yes |
| | | MBNL3 | 0.8 | Yes |
| | | GM2A | 0.32 | Yes |
| | | TXK | 0.28 | No |
| | | FAM46A | 0.24 | No |
| | | MYLIP | 0.2 | No |

Table S4: Existing knockoff generating method for benchmark in this paper

| Type | Subtype | Details | Limitations | Type limitation | In Benchmark |
|---|---|---|---|---|---|
| Second-order Approximation | SDP | oracle (simulation), JS-shrunk | Low power | Fail when $P_X$ contain higher orders; Affected by covarince estimation | T |
| | MVR | | Not implemented in R | | T |
| | Entropy Knockoff (ME) | | Computation is slow in R | | T |
| | Deep Knockoff | | Large sample size required | Large sample size required | F |
| SCIP (Approximate) | KnockoffScreen | K-nearest neighbour; linear regression | OLS is not often valid; K is too small in high-dimension | Trade-off in speed and accuracy | F |
| | SeqKnockoff | lasso or elastic net regression | Computationally very slow | | T |
| | PCKO | PC-regression | Computationally slow | | T |
| Structured correlation | IPAD | | Assume X follows factor model | Might fail when X not follow factor model | T |

$$T = \min \left\{ t > 0 : \frac{\#\{j : W_j \leq -t\}}{\#\{j : W_j \geq t\} \vee 1} \leq q \right\}.$$

Therefore, the false discovery proportion (FDP) of set $\hat{S}$ can be controlled, as follows:

$$\mathrm{FDP}(t) \approx \frac{\#\{j \in \mathcal{H}_0 : W_j \geq t\}}{\#\{j : W_j \geq t\}} \approx \frac{\#\{j \in \mathcal{H}_0 : W_j \leq -t\}}{\#\{j : W_j \geq t\}} \leq \frac{\#\{j : W_j \leq -t\}}{\#\{j : W_j \geq t\}} \leq q,$$

and the procedure of selecting controls the modified FDR defined as

$$\mathrm{mFDR} = \mathbb{E}\left[ \frac{\#\{j : j \in \hat{S} \cap \mathcal{H}_0\}}{|\hat{S}| + 1/q} \right].$$

Slightly more conservatively, given by incrementing the number of negatives in discoveries by 1, knockoff+ defines the threshold as

$$T_+ = \min \left\{ t > 0 : \frac{1 + \#\{j : W_j \leq -t\}}{\#\{j : W_j \geq t\} \vee 1} \leq q \right\}.$$

Selecting $\hat{S} = \{j : W_j \geq T_+\}$ controls the usual FDR as,

$$\mathrm{FDR} = \mathbb{E}\left[ \frac{\#\{j : j \in \hat{S} \cap \mathcal{H}_0\}}{|\hat{S}|} \right].$$

**Algorithm S1** Sequential Conditional Independent Pairs (SCIP), Candès et al. (2018)

$j = 1$
**while** $j \leq p$ **do**
  Sample $\widetilde{X}_j$ from $\mathcal{L}(X_j | X_{-j}, \widetilde{X}_{1:j-1})$, conditionally independently from $X_j$
  $j = j + 1$
**end while**

#### S3.2 Review of existing knockoff generators

There are many possible ways to construct valid knockoff variables. Although it is theoretically ensured that knockoff variables with model-X properties controlled FDR, the power of knockoff filtering varies among different construction approaches. An ideal knockoff construction method can identify as many variables as possible (i.e. maximum power) while controlling the FDR under the desired level. Two main groups of knockoff-generating approaches are described below (Summarised in Table S4).

##### S3.2.1 Joint distribution approximation knockoff generators

The first type of knockoff variable generators are based on the approximation of the property 1 of model-X. For example, $\widetilde{X}$ is a second-order knockoff copy when the first two orders of joint distribution $(X, \widetilde{X})$ and $(X, \widetilde{X})_{swap(S)}$ are matched. This condition is equivalent to $\mathbb{E}(X) = \mathbb{E}(\widetilde{X})$ and

$$\operatorname{cov}(X, \widetilde{X}) = \begin{bmatrix} \Sigma & \Sigma - \operatorname{diag}\{s\} \\ \Sigma - \operatorname{diag}\{s\} & \Sigma \end{bmatrix}, \tag{2}$$

where $\Sigma$ is the covariance matrix of $X$ and $s \in \mathbb{R}^p$ such that 2 is positive semidefinite. Then, to improve the power of second-order approximation knockoffs, different algorithms might be applied to solve $s$. In the original model-X paper (Candès et al., 2018), semidefinite programming (SDP) tried to maximise the power by minimising the marginal correlation between the knockoff variable and its original variable. However, this method may show very low power in some settings, such as high-correlated or high-dimension settings. This can be observed in our simulation studies (e.g. Figure S2). Minimised reconstructability (MRC) knockoff (Spector and Janson, 2022) proposed another $s$-solving algorithm that maximises the conditional variance of variables conditioning on the other variables and knockoff variables or maximises the entropy of the joint distribution of $X$ and $\widetilde{X}$. Overall, second-order approximation knockoff construction requires mean and covariance of $X$ known or estimated and is the exact construction for $X$ following multivariate Gaussian and second-order approximation for other distributions. When the covariate distribution does not follow multivariate Gaussian, the second-order approximation knockoff might fail to control FDR, especially when the relationship among covariate $X$ is non-linear and cannot be fully described by the first two orders. This limitation is shown in the simulation Section **??** and Figure **??**.

Deep knockoffs (Romano et al., 2020) can generate knockoff variables also based on the joint distribution approximation but with higher-order matching. Similar to other deep learning models, this knockoff construction method also requires a very large sample size to train.

##### S3.2.2 SCIP knockoff generators

A general algorithm for the exact construction of the knockoff variable has been proposed for any type of $X$ (Candès et al., 2018), namely sequential conditional independent pairs (SCIP). The SCIP algorithm (Algorithm SS1) is based on the derived property of model-X knockoffs that any pair $(X_j, \widetilde{X}_j)$ is exchangeable conditioning on all other variables and their knockoffs so that for each $j \in (1, \cdots, p)$, $\widetilde{X}_j$ can be generated by sampling from $\mathcal{L}(X_j|X_{-j}, \widetilde{X}_{1:j-1})$. However, implementing this algorithm is complicated or not always practical, since the conditional distribution has to be recomputed at each step. Exact sampling methods have been implemented for Gaussian distributions or distributions where the latent variables can be represented as a Bayesian network (BN), such as Sesia et al. (2019) for discrete and hidden Markov chain models with application in GWAS and Gimenez et al. (2019) for Gaussian mixture models. This proposed knockoff sampling algorithm relaxes the conditional distribution $\mathcal{L}(X_j|X_{-j}, \widetilde{X}_{1:j-1})$ to only condition on a subset, that is, neighbours of $X_j$ on the Bayesian network, instead of all other variables and constructed knockoffs, thus reducing the computational burden.

In addition to exact SCIP sampling (e.g., for the Markov chain model in GWAS), several approximate SCIP-based methods have been proposed and applied in more general biological studies without any distribution assumption on $X$. Generally, these approximate sampling methods use different approaches to approximate the conditional distribution, i.e., $\mathcal{L}(X_j|X_{-j}, \widetilde{X}_{1:j-1})$, sequentially for each variable. For example, Sequential Knockoffs, proposed by (Kormaksson et al., 2021), use prediction residuals from lasso or elastic net (EN) as the conditional distribution. `KnockoffScreen` for GWAS (He et al., 2021) used linear regression with the most correlated variables as predictors. Jiang et al. (2021) used principal component (PC) regression in their knockoff boosted tree (`KOBT`) model to sequentially fit the dependence relationship in each variable with the other $j - 1$ variables. These approximate SCIP-based knockoff construction methods showed empirically valid in most of the cases provided in their literature. However, there are limitations to their applications in more challenging cases, such as high-dimension studies. Sequential knockoffs can be computationally slow when $X$ consists of hundreds of variables, since lasso and EN are iteratively fitted with cross-validation for every variable in the data set; OLS in `KnockoffScreen` is not always feasible because of the singularity when $X$ is high-dimensional or high-correlated; PC-regression-based SCIP knockoff generation method, `KOBT`, might be not able to capture the dependence among variables sufficiently with a low number of PCs (leading to inflated FDRs) or might capture redundant variance with a large number of PCs (leading to low powers) since PCs in PC regression are the eigenvectors with the largest eigenvalues that capture most of the total variance in the other $j - 1$ variables instead of capturing the variance that can explain the most of variance of $X_j$.

###### S3.2.3 IPAD

Intertwined probabilistic factors decoupling (IPAD) (Fan et al., 2020) assumes $X$ follows the exact factor model, i.e., $X = X = TP' + E = C + E$, and estimate parameters $(\theta)$ of $C$ and $E$. Then the knockoffs can be construct as $\widetilde{X}(\hat{\theta}) = \hat{C} + \widetilde{E}$, where $\widetilde{E}$ is independently sampled from estimated distribution of $E$. In the IPAD paper, the low-rank structure $C$ is estimated by SVD and the residual $E$ is estimated as an i.i.d. centred normal distribution with variance equal to the empirical variance of the residual.

#### S3.3 PLSKO

Here we proposed a new knockoff variable construction method, 'PLSKO', aiming to control the FDR effectively in high-dimensional biological data and meanwhile to reach as high power as possible. PLSKO is an approximate sequential conditional independent pairs (SCIP) approach Candès et al. (2018) (described in Algorithm S1 and Figure **??**).

The PLSKO procedure to generate knockoff variable $\widetilde{X}$ for a dataset $X \in \mathbb{R}^{n\times p}$ is described below (also see Algorithm S2):

- *Step 1: Generating neighbour set for each variable $X_j, j = 1, \ldots, p$.*
  To lower the computational burden and inspired by Gimenez et al. (2019), Sesia et al. (2019) and He et al. (2021), the conditioning distribution $\mathcal{L}(X_j|X_{-j}, \widetilde{X}_{1:j-1})$ can then go down to $\mathcal{L}(X_j|X_{k\in BN_j}, \widetilde{X}_{1\le k\le j-1, k\in BN_j})$, where $BN_j$ is the neighbours of $X_j$ in the Bayesian network of $X$. Instead of controlling all the other variables $(X_{-j})$ and knockoff variables before $(\widetilde{X}_{1:j-1})$, now the conditional distribution is only on those in the neighbourhood of $X_j$, and other variables are assumed to be independent with $X_j$. When there is no user-prespecified neighbour list, PLSKO uses the sample correlation with $X_j$ as the similarity measure and defines the nearest variables either with a correlation greater than a user-defined absolute correlation threshold or with a correlation in the top quantile of all pairwise correlations in the sample. The choice of absolute or quantile threshold is to ensure that $P(X_j|X_{k\in BN_j}, \widetilde{X}_{1\le k\le j-1, k\in BN_j})$ accurately mimics $P(X_j|X_{-j}, \widetilde{X}_{1:j-1})$ and to avoid overfitting.

- *Step 2: Fit $X_j$ as a function of $X_{k\in BN_j}$ and $\widetilde{X}_{1\le k\le j-1, k\in BN_j}$.*
  To generate knockoff variables from $\mathcal{L}(X_j|X_{k\in BN_j}, \widetilde{X}_{1\le k\le j-1, k\in BN_j})$, we assume a model,

  $$X_j = g(X_{k\in BN_j}, \widetilde{X}_{1\le k\le j-1, k\in BN_j}) + \epsilon_j,$$

  where $\epsilon_j$ is a random error term. In other words, $g(\cdot)$ is the component of the information in $X_j$ that can be explained/represented by the other variables and constructed knockoff variables; $\epsilon_j$ is the unique and irrepretentalbe information in $X_j$. PLSKO approximates $g(\cdot)$ as a linear model, that is, $g(X_{ij}|X_{k\in BN_j}, \widetilde{X}_{1\le k\le j-1, k\in BN_j}) = \alpha + \sum_{k\neq j, k\in BN_j} \beta_k X_{ik} + \sum_{k\le j-1, k\in BN_j} \gamma_k \widetilde{X}_{jk}$, and uses partial least squares regression (PLS regression, described in Appendix section S3.3.2) to estimate parameters in model $g(\cdot)$ and calculate the fitted value of $X_j$, namely $\hat{X}_j$.

- *Step 3: Generate knockoffs by permuting the residuals within samples.*
  Residuals are then calculated as $\hat{\epsilon_j} = X_j - \hat{X}_j$, as an approximation of the conditional distribution $\mathcal{L}(X_j|X_{-j}, \widetilde{X}_{1:j-1})$, representing the part of $X_j$ that cannot be predicted by the other variables and the generated knockoff variables. Intuitively, we see the predictable part, $\hat{X}_j$, as the part that needs to be controlled and the part that preserves the dependency relationship among the variables, thus is to be preserved in knockoff variables as well. We see the residual as the unpresentable unique information of $X_j$, independent with the residuals of other variables. Only if the residual is correlated with the outcome, we could say (but not always without other conditions) $X_j$ might be important as it is not independent of the outcome $y$ conditioning other variables. In knockoff construction in PLSKO, we generate knockoff variable $\widetilde{X_j}$ by adding up $\hat{X}_j$ and the permuted residual $\widetilde{\epsilon_j}$. Permutation of residuals instead of sampling from a specified and parameterised conditional distribution avoids more assumptions and allows PLSKO to be applied on data from more types of distribution.

**Algorithm S2** Partial Least Square Knockoff (PLSKO)

**Require:** Dataset $X \in \mathbb{R}^{n \times p}$; Neighbour set for each $X_j$: $BN_j,\ j = 1, \ldots, p$

Number of components $r$ in PLS regression

Sparsity applied to sparse PLS regression $s$

$j = 1$

Centered $X$ with $X - \mu$, $\mu \in \mathbb{R}^p$ is the sample mean of $X$

**while** $j \leq p$ **do**

Define the neighbour knockoff set of $X_j$: $BN_{ko,j} = \{k | k \in BN_j$ and $k \leq j - 1\}$

**if** $|BN(X_j)| = 0$ **then**

$\hat{X}_j \leftarrow 0$

**else if** $|BN(X_j)| = 1$ **then**

Fit OLS regression as $y_{OLS} = X_j$, $X_{OLS} = [X_{k \in BN_j}, \widetilde{X}_{BN_{ko,j}}]$ (augmented matrix)

Calculate $\hat{X}_j$

**else**

Fit PLS regression as $Y_{PLS} = X_j$, $X_{PLS} = [X_{k \in BN_j}, \widetilde{X}_{BN_{ko,j}}]$ (augmented matrix), with $r$ components;

Calculate $\hat{X}_j = \hat{X}_{j,PLSreg}$ on the first $r$ components

**end if**

Get $\tilde{\epsilon}_j$ by permuting the residual $\hat{\epsilon} = X_j - \hat{X}_j$;

Calculate $\widetilde{X}_j = \hat{X}_j + \tilde{\epsilon}_j$;

$j = j + 1$

**end while**

$\widetilde{X} = \widetilde{X} + \mu$

**Output:** $\widetilde{X}$

###### S3.3.1 PLS-AKO

In case studies, we generated multiple knockoff variables for each observed fixed dataset and incorportated by aggragation of multiple knockoff (AKO) (Nguyen et al., 2020) to accommodate for the randomness and improve the stability from knockoff variable generation. To distinguish between PLSKO with a single run and multiple PLSKO incorportated by AKO, we refer the later one as 'PLS-AKO'. The PLS-AKO used in this study is briefly described as the following steps:

- **Step 1: Construct multiple knockoff by PLSKO.** $B$ knockoff variables are constructed by PLSKO parallel as $\widetilde{X^{(b)}} \in \mathbb{R}^{n\times p}$, $b = 1, \ldots, B$.

- **Step 2: Calculate the importance statistics.** $X$ and each constructed knockoff variable set $\widetilde{X^{(b)}}$ are run into the (logistic) lasso regression, and the LCD is used as the importance statistics $W^{(b)} \in \mathbb{R}^p, b = 1, \ldots, B$.

- **Step 3: Calculate intermediate p-value.** For each $W^{(b)}$, calculate the empirical $p-$value for $X_j, j = 1, \ldots, p$ as:

$$\pi_j^{(b)} = \begin{cases} \frac{1+\#\{k:W_k\leq -W_j\}}{p} & \text{if } W_j > 0 \\ 1 & \text{if } W_j \leq 0 \end{cases} \tag{3}$$

- **Step 4: Aggregate empirical $p-$value.** For each $X_j$, $B$ empirical $p-$values, $\pi_j^{(b)}$, are aggregated by the quantile aggregation procedure as:

$$\tilde{\pi}_j = \min\left\{1, \frac{q_\gamma\left(\{\pi_j^{(b)} : b \in [B]\}\right)}{\gamma}\right\} \tag{4}$$

  where $q_\gamma(\cdot)$ is the $\gamma$-quantile function. As tested and recommended in Nguyen et al. (2020), we fix $\gamma = 0.3$, and $B = 50$ as 50 knockoff variables has been generated.

- **Step 5: Adjust $p-$value to control the FDR.** With $\tilde{\pi} \in \mathbb{R}^p$, the Benjamini-Hochberg (BH) procedure is used to control the FDR. Variables with the $\tilde{\pi}$ less or equal than the BH threshold are selected by PLS-AKO.

In practice, we found that PLS-AKO generally had lower FDR and higher power compared to the average of PLSKO before aggregation.

###### S3.3.2 Review of Partial Least Square (PLS) regression

Partial Least Squares (PLS) (Wold, 1966) is a wide class of techniques for modelling relations between sets of observed variables by means of latent variables, assuming that observed data, including predictors and multiple responses, are all driven by a small number of latent variables and the latent variables from the two datasets are highly-related. PLS has been widely applied to high-dimensional biological data for a variety of research questions such as classification and biomarker identification, due to the ability of PLS to work very well for data with very small sample sizes and a large number of parameters and high computational and statistical efficiency (Boulesteix and Strimmer, 2007).

The general underlying model of multivariate PLS is: consider predictor $X \in \mathbb{R}^{n\times p}$ and response $Y \in \mathbb{R}^{n\times q}$, we assume both of them follow the factor model, that is:

$$\begin{aligned} X &= TP^\top + E \\ Y &= UQ^\top + F, \end{aligned} \tag{5}$$

where the $T$and $U \in \mathbb{R}^{n\times r}$ are matrices of the $r$ extracted score vectors (components, latent factors), the matrix $P \in \mathbb{R}^{p\times r}$ and $Q \in \mathbb{R}^{q\times r}$ are the matrices of loadings, and $E \in \mathbb{R}^{n\times p}$ and $F \in \mathbb{R}^{n\times q}$ are the matrices of residuals, assumed to be independent and identically distributed random normal variables. The decompositions of $X$ and $Y$ are made so as to maximise the covariance between $T$ and $U$, i.e., iteratively find vectors $w, c$, such that $\arg\max_{\|w\|=1,\|c\|=1} w^\top X^\top Y^\top c$, and then $t = Xw, u = Yc$.

PLS is an iterative process. After the extraction of the score vectors $t$, $u$ the matrices $X$ and $Y$ are deflated by subtracting their rank-one approximations based on $t$ and $u$. Different forms of deflation define several variants of PLS (Rosipal and Krämer, 2006). For example, canonical PLS (Mode A) deflates both $X$ and $Y$ in each iteration. The relation between the two blocks is symmetric in this mode, and it is more appropriate for modelling existing relations between sets of variables (e.g., investigating relationships between RNA and proteins) in contrast to prediction purposes. In contrast, PLS regression (Höskuldsson, 1988) assumes $T = \{t_i\}_{i=1}^r$ (the latent variables of $X$) are good predictors of $U$ (the latent variables of $Y$). Deflation in regression mode is to remove a component of the regression of $Y$ on $t$ (the latent variables of $X$, instead of $u$) from $Y$ at each iteration.

**PLS regression** To introduce our new PLSKO method, we briefly describe the steps of PLS regression and prediction. When predicting $Y$ from $X$, PLS regression assumes that $T$ are good predictors of $U$, i.e., $U = TD + H$, where $D \in \mathbb{R}^{p\times p}$ is a diagonal matrix, so the equation 5 can be written as:

$$Y = UQ^\top + F = (TD + H)Q^\top + F = TDQ^\top + (HQ^\top + F) = TC^\top + F^*, \tag{6}$$

where $C^\top = DQ^\top \in \mathbb{R}^{n\times q}$ is the regression coefficients and $F^* = HQ^\top + F$ is the residual matrix. Again, from equation 5, we have $T = XW(P^\top W)^{-1}$, so when fitting $Y = XB + F^*$, the regression coefficients $B = W(P^\top W)^{-1}C^\top = X^\top U(T^\top XX^\top U)^{-1}T^\top Y$. For a new set of $X_{new}$,

$$\hat{Y}_{PLSreg} = X_{new}\hat{B} = X_{new}X^\top U(T^\top XX^\top U)^{-1}T^\top Y. \tag{7}$$

In this paper, PLS regression is calculated by Non-linear iterative partial least-squares (NIPALS) algorithm (Wold, 1975).

**Advantages of PLS regression for SCIP** In high-dimensional data, where $n < p$, or in highly correlated data, where the rank of the data $r < p$, the usual linear regression with ordinary least squares (OLS) cannot be applied, since the covariance matrix of $X$ (which can have a maximum rank $n - 1$) is singular. That is, OLS cannot be applied to conditional distribution calculation in the SCIP algorithm when neighbour variables and their knockoffs outnumber the sample size. Otherwise, reducing the neighbour numbers lower than the sample size by placing a sparser neighbour filter might lead to incorrect conditional distribution calculation due to insufficient covariates, then the violation of the exchangability requirement of the model-X knockoff, and thus inflated FDR. In contrast, PLS regression can be applied to cases in which $n < p$ and $r < p$ without reducing the number of predictors. Principal component regression (PC regression) is also a reduced rank regression method, which firstly extracts the first components (called principal components) that capture most of the total variance and then uses the first principal components in the data as the predictor variables in the regression. Principal component extraction in PC regression does not use the response for component construction. In contrast, PLS regression takes the response variable $y$ into account for the construction of the components. This feature allows PLS regression to capture more information in the response variable $y$ with a lower number of components than PC regression and better performance in prediction problems. Moreover, in PLS regression, dimension reduction and regression are performed simultaneously. As the SCIP algorithm is iterative on each variable in the dataset, the computational efficiency for conditional distribution is high and therefore quite advantageous for high-dimensional data. The lower number of components and the simultaneous fitting make PLS regression more efficient than PC regression. In summary, the use of PLS regression in the computation of the conditional distribution ensures accuracy with enhanced efficiency.

#### S3.4 Benchmark methods

In the simulation experiments in this study, second-order approximation (SOA) knockoff generators for benchmark include: (they may categorised by the algorithms for solving $s$ and the covariance estimators)

- *oracle*: Semidefinite programming (SDP) with the known oracle covariance matrix, implemented by the `R` package `knockoff`;
- *JS*: SDP with the covariance estimated by James-Stein-type Shrinkage (JS) (Ledoit and Wolf, 2003) (JS), implemented by the `R` package `knockoff` and `corpcor`;
- *mvr-oracle*: Minimised the variances-based reconstructability (MVR) algorithm with the oracle covariance, implemented by the `Julia` package `Knockoffs`;
- *mvr-JS*:MVR with the JS-type estimated covariance, implemented by the `Julia` package `Knockoffs`;

- *maxent-oracle*: Maximised the entropy (MAXENT) algorithm with the oracle covariance, implemented by the `Julia` package `Knockoffs`; (An `R` alternative can be found in the package `cheapknockoff` (Yu et al., 2022), but could be much slower.)
- *maxent-JS*: MAXENT with the JS-type covariance estimator, implemented by the `Julia` package `Knockoffs`;
- *maxent-approx*: MAXENT with the covariance estimated as a block diagonal structure with `windowsize = 100`, implemented by the `Julia` package `Knockoffs`.

In the simulation experiments in this study, SCIP knockoff generators for benchmark include:

- *PC*/*PCKO* (in simulation): SCIP with prior neighbourhood selection and with PC regression as the conditional distribution approximation. Same parameter settings (i.e., number of components and neighbour threshold) used with PLSKO;
- textitPCKO(in semi-simulation): SCIP without neighbour selection and 'full' components, with PC regression as the conditional distribution approximation, implemented by the `R` package `KOBT`. Same parameter setting with PLSKO-full.
- *seqko*: SCIP with lasso or elastic net as the conditional distribution approximation, implemented by the `R` package `seqknockoff` (https://github.com/kormama1/seqknockoff).

In this paper, the numbers of components used in IPAD were the true rank of simulation data, or determined by the $PC_{p1}$ criterion in the semi-simulation study, same as PLSKO-full.

#### S3.5 Simulation studies

Below we describe in detail the simulation of a single data set $(X, y)$ for a given parameter configuration. For each parameter configuration we simulate $n_{sim} = 100$ data sets and apply the knockoff filtering with different knockoff variable generating methods.

##### S3.5.1 Data generation

- **Simulation of $X$.** We simulate the $n \times p$ design matrix $X$ from a block factor models. Each block of $X$ is generated from $X_b = F_b\Lambda_b' + E_b$, where latent factors $F_b \in \mathbb{R}^{n\times r_b}$ the factor loadings $\Lambda_b \in \mathbb{R}^{p_b \times r_b}$, and $E \in \mathbb{R}^{n\times p_b}$ $\sim \mathcal{N}(0, I_{p_b})$. The correlation structure of $X$ from the same block is represented by the low-rank factor component $F_b\Lambda_b'$ and is expected to be controlled in the knockoff filter. Variables from different blocks are independent. We set 5 blocks and the number of variables in each block as $p_b = 100$ and the number of latent factors as $r_b = 3$. Factor loadings are sampled independently from a uniform distribution $U(1, 2)$ with a random sign of equal probability.
- **Simulation of $X$ for robustness tests.** We simulate $X$ from a block multivariate Gaussian and block quadratic factor model. The blocks of multivariate normal $X$ are generated from $\mathcal{N}(\mathbf{0}, \mathbf{\Sigma_{p_b}})$, with $\Sigma = (\sigma_{ij})$, $\sigma_{ij} = \rho^{|i-j|}$ (autoregressive process of order one, AR1) or $\sigma_{ij} = \rho$ (equicorrelation setting). $\rho$ is set to be 0.5. In each block of the quadratic factor model, half of the $p_b$ variables are generated from the factor model (as described above) and the other half of the $p_b$ variables are generated as the square of the variables from the generated half. This quadratic model provides a scenario in which the dependency among variables is non-linear, i.e., $X$ is not sufficiently described by the first two moments.
- **Simulation of $y|X$.** The response variable $y$ is simulated from $y_i = f(x_i) + (c \times s \times p)\epsilon_i$. $c$ is a constant controlling the signal-to-noise ratio, $s \times p$ is the number of true signal variables, and *epsilon* is the model error following $\mathcal{N}(0, 1)$. $X$ is rescaled column-wise before generating $y$. Without further specification, $c = 0$ and $s = 0.1$. For continuous $y$, $f(x) = x\beta$, where $\beta$ is the coefficients sampled from $U(3, 5)$ with a sign following Bin(0.5) for the signal variables and $\beta = 0$ for the null variables; for the categorical $y$, $f(x) = \text{Bin}(\pi(x\beta))$, where $\pi$ is the Sigmoid function.

Table S5: Summary of configurations in simulation experiments

| Experiment | $X$ | | | | | | $y$ | | | Report in |
|---|---|---|---|---|---|---|---|---|---|---|
| | Distribtution | Number of latent factor per block, $r$ | Correlation, $\rho$ | Sample size, $n$ | Blocks, $b$ | Variables in each block, $p_b$ | Type | Sparsity, $s$ | Noise, $c$ | |
| Varying sample size | Factor model | 3 | NA | 150-250 | | | linear | 0.1 | 0 | Figure S2 |
| Varying sparity | Factor model | 3 | NA | 500 | | | linear | 0.1 - 0.4 | 0 | Figure S3a |
| Varying sigal-noise ratio | Factor model | 3 | NA | 500 | 5 | 100 | linear | 0.1 | 1:16- 5:16 | Figure S3b |
| Robustness test | Multivariate Gaussian: AR1 and Equicorr | NA | 0.5 | 150-250 | | | linear | 0.1 | 0 | Figure S4b and S4a |
| Robustness test | Quadratic factor model | 3 | NA | 150-500 | | | linear | 0.1 | 0 | Section **??** (Figure **??**) |
| Binary $y$ | Factor model | 3 | NA | 200-1000 | | | binary | 0.1 | 0 | Figure S5 |

###### S3.5.2 Simulations setting

- **Varying sample size** In this experiment we vary the sample size $n \in \{150, 160, \ldots, 250\}$ and compare the FDP and TPP across the benchmark methods.
- **Varying sparsity** We vary the sparsity $s \in \{0.1, 0.2, 0.3, 0.4\}$ (i.e., 50, 100, 150, 200 important variables are used to generate $y$) with fixed sample size $n = 500$.
- **Varying signal-noise ratio** We vary the signal-noise ratio $c \in \{1, 2, \ldots, 5\}$, corresponding to the approximate ratio of unmeasured important variables to the measured important variables $1:16, 1:8, \ldots, 5:16$ with fixed sample size $n = 500$.
- **Robustness test with varying sample size** We vary the sample size $n \in \{150, 160, \ldots, 250\}$ in two multivariate gaussian design $X$ and quadratic factor model to examine the robustness of PLSKO and other methods in different data distribution.
- **Binary response variable** $y$ We generate the binary response $y$ by applying the sigmoid function on the centred linear combination of randomly selected important variables, and test on varying the sample size from 200 to 1000. Due to the near-zero power when controlling on the usual FDR (with $T_+$ threshold), we only control the modified FDR.

A summary is available in Table S5.

###### S3.5.3 Performance measures

We use the false discovery proportion (FDP) and the true positive proportion (TPP) as the measure of performance for the knockoff generators, which are defined as:

$$\mathrm{FDP} = \frac{V}{R} \tag{8}$$

where $V$ is the number of false positives in the selected variable set and $R$ is the total number of rejected hypotheses (selected variables), and

$$\mathrm{TPP} = \frac{S}{m_1} \tag{9}$$

where $S$ is the number of true positives in the selected variable set and $m_1$ is the total number of true alternative hypotheses (i.e., the known important variables).

As mentioned in Section **??**, FDR of knockoff filtering with different knockoff generating methods are estimated as the average FDP and power are estimated as the average TPP over the repeated experiments.

A good knockoff variable generating method should control FDR under the nominal level and achieve as high power as possible.

#### S3.6 Preeclampsia datasets

###### S3.6.1 Data description

A summary of the data used for semi-simulation and the case studies analyses is presented in Table S6.

Table S6: Summary of data used in semi-simulation and case study.

| Data | Samples | Type | Predictors | Reseponse | Used in |
|---|---|---|---|---|---|
| Moufarrej et al. (2022) Circulating cell-free RNA for preeclampsia prediction (GSE192902) | n = 71 (16 preeclamptic, 55 normotensive) | Trascriptomics: RNA-seq | 7160 genes | Synthetic | Semi-simulation in Section ??; Figure ?? |
| | | | 200 most variable genes | Disease condition | Case Study in Section ??; Table S2 |
| | | | 81 genes elevated in placenta | Disease condition | Case Study in Section ??; Table ?? |
| | | | | RNF14 gene expression level | Case Study in Section ??; Table S3 |
| Marić et al. (2022) Multiomics for preeclampsia prediction | n = 36 (18 preeclamptic, 18 normotensive); n = 49 (29 preeclamptic, 20 normotensive, only microbiome ) | Proteomics | 1303 proteins | Synthetic | Semi-simulation in Section ??; Figure S7a,S6b,S7b |
| | | Urine Metabolics | 8171 metabolites | | |
| | | Microbiomics 16s rRNA | 144 OTUs | | |
| | | Plasma Transcriptomics: RNA-seq | 74 genes elevated in placenta | Disease condition | Case Study in Section ??; Table ?? and S1 |
| | | Proteomics | 36 proteins released by placenta | | |
| | | Microbiomics 16s rRNA | 144 OTUs | | |

**Cell-free RNA sequence data in human serum with preeclampsia** The dataset contains circulating cell-free RNA-seq (cfRNA) data previously collected as part of a prospective longitudinal study of 71 pregnant women (19 with preeclampsia, 52 normotensive controls) by Moufarrej et al. (2022). The filtered count matrix was obtained from GSE192902, including 7,160 genes with a level of at least 0.5 count-per-million (CPM) in at least 75% of samples. The details of the study design, the sample collection, the cfRNA library preparation and data quality assessment were previously described in Moufarrej et al. (2022). We only used samples that were sampled before 12 weeks of gestation (sampling time 1) to avoid repeated measurements in the same individual in our analysis, given the assumption of *i.i.d.* in the knockoff filtering.

**Multi-omics study in human with preeclampsia** To test the robustness of knockoff generating methods in different types of data, we use data from a multi-omics longitudinal study for preeclampsia of 36 pregnant women (18 with preeclampsia and 18 normotensive controls) from cohort 1 or 49 women (29 preeclampsia, 20 normotensive controls, microbiome only) from both cohorts (Marić et al., 2022). The datasets include transcriptomic, metabolic, proteomic data from serum, metabolic data from urine and microbiome data from virginal swap which were sampled from the same individual at the same time point, consisting of 37184 genes, 3621, 1305, 8717 and 1255 variables. Same as the cfRNA-seq data, we only use samples from the first cohort and the first time point (generally before 12 weeks of gestation) in our analysis. Variable prefiltering is applied to the transcriptome data and microbiome data. In transcriptomics data, 6394 genes with a level of at least 0.5 count-per-million (CPM) in at least 75% of samples are kept for the following analysis. In microbiome data, only one OTU from each almost identical pair (sample correlation $>0.95$) is kept. Then, we remove OTUs for which the sum of counts is below a set threshold (0.01%) compared to the total sum of all counts, leaving 144 OTUs kept for the following analysis. For the comparison with *limma* in the case study, we also normalised the microbiome count matrix with TMM.

##### S3.6.2 Data preprocessing

For transcriptome data (RNA-seq), Trimmed mean of M-values (TMM) and Counts per million (CPM) are used for normalisation to account for differences in library sizes potentially caused by different sequence depth between samples. Imputed and normalised proteome and metabolome data are log-2 transformed (described in Marić et al. (2022)). Log-ratio transformation with Centered Log Ratio (CLR) was applied to microbime data. Taxonomic assignment was performed against the Silva v138.1 database using the `R` package dada2 (Callahan et al., 2016).

#### S3.7 Semi-simulation: real $X$ with synthetic $y$

Given the validity of knockoff generators only relies on the distribution of $X$ (see the model-X knockoff definition in Section ??), we check the robustness of knockoff generators on real biological data by simulating artificial response data using the real covariate data. The true important variable sets in this semi-simulation are still known, so we are able to assess the FDPs and TPPs for knockoff generators.

According to the results of simulation experiments where $y$ is linearly dependent on $X$, with the fixed sample sizes of the real datasets, we set up the number of variables ($p$) for semi-simulation experiments with the $n : p$ ratio ranging from 5:5 to 2:5 in order to keep the TPPs above zero. For the cfRNA dataset with a sample size of 71, we run the semi-simulation with $p = 100, 150, 200$; for the multi-omics dataset with a sample size of 36, we run with $p = 40, 80, 120$. For each iteration, a subset of variables is randomly selected as the dataset for testing to mimic the distribution of the real data. Artificial important variables (25 for scRNA data and 15 for multi-omics) are randomly selected and used for generating the artificial response $y$ by linear combination. Knockoff filtering with different knockoff-generating methods for the benchmark is then applied with lasso LCD as importance statistics and the more liberal threshold $T$ to control the modified FDR (mFDR) (Candès et al., 2018). Selected variables are then compared with the known true variables to calculate the FDP and TPP. For each real dataset, semi-simulation experiments are run 50 times with the target FDR at 0.05 for cfRNA data or 0.10 for datasets from the multi-omics study.

#### S3.8 Case studies

##### S3.8.1 Subsets of variables

We further lowered the number of variables before applying knockoff filtering given the observed limitation of near-zero power when the $p : n$ ratio is too large. For the cfRNA data, we selected 200 genes with the highest variance and apply PLSKO and knockoff filtering to identify variables associated with preeclampsia. Semi-simulation is also run to confirm the validity.

Alternatively, we also reduced variable size according to biological knowledge, leading to more specific biological questions. As cfRNA is derived from many tissues in the body, we also select genes that are placental-specific elevated measured in the HPA(v.19; https://v19.proteinatlas.org/humanproteome/tissue/placenta; or the most updated v23, results showed in Supplementary)(Uhlen et al., 2019), aiming to identify physiological changes associated with preeclampsia in the first-trimester placenta. Eighty-one placental-specific elevated genes are found in the prefiltered cfRNA dataset. Similarly, 36 placenta-derived proteins were kept in the proteomics data from the multi-omics study, according to the identification study by Degnes et al. (2022).

##### S3.8.2 Application of knockoff filtering

For each dataset, knockoff filtering with the difference of coefficients in the logistics lasso regression as the importance statistics was run 50 times with the target FDR level 0.05. The more liberal threshold $T$ to control the modified FDR (mFDR) (Candès et al., 2018) was used for higher power. Selection frequency of the variables is reported when their selection frequency higher than 0.1. Aggregation of multiple knockoffs (AKO, Nguyen et al. (2020)) was then used to integrate the repeatedly generated knockoff variables to improve the stability with $\gamma = 0.3$, namely PLS-AKO.

#### S3.9 Software packages used

All the experiments were run in `R` (v 4.3.0) and `Julia` (v 1.9.3), with following listed packages:

- **mixOmics** `R` package (Rohart et al., 2017) (v 6.25.1): used for PLS regression and sparse PLS regression.
- **knockoff** `R` package (Patterson and Sesia, 2022) (v 0.3.6): used for the second-order approximation knockoff construction with SDP and generate LCD.
- **glmnet** `R` (Friedman et al., 2010; Tay et al., 2023) package (v 4.1-8) : used to generate LCD.
- **RSpectra** `R` package (v 0.16-1): used for IPAD knockoff and PCKO.
- **JuliaCall** `R` package (v 0.17.5): used to call Julia package in R environment.

- **Knockoffs** `Julia` package (Chu et al., 2024) (v 1.1.5): used for the second-order approximation knockoff construction with MVR and ME.
- **Grace-AKO** `R` package (Tian et al., 2022): used for knockoff aggregation (AKO).
- **edgeR** `R` package (v 3.42.4) (Robinson et al., 2010): used for real data preprocess.
- **limma** `R` package (v 3.56.2) (Smyth, 2005): used for case studies.
- **doParallel** `R` package (v 1.0.17): used for parallel computing in simulation and generating multiple knockoffs in case studies.
- **tidyverse** `R` package (v 2.0.0).
- **Dada2** `R` package (v 1.28.0) Callahan et al. (2016): used for microbiome data taxonomic assignment.

Herman Wold. Path models with latent variables: The NIPALS approach., 1975.

Guo Yu, Daniela Witten, and Jacob Bien. Controlling costs: Feature selection on a budget. Stat, 11(1):e427, 2022. ISSN 2049-1573. doi:10.1002/sta4.427. URL https://onlinelibrary.wiley.com/doi/abs/10.1002/sta4.427.
